# Critical Assessment of MetaProteome Investigation 2 (CAMPI-2): Multi-laboratory assessment of sample processing methods to stabilize fecal microbiome for functional analysis

**DOI:** 10.1101/2025.01.31.635836

**Authors:** Alessandro Tanca, Kay Schallert, Lucia Grenga, Samantha L. Peters, Marcello Abbondio, Laura De Diego, Maria Antonietta Deledda, Sven-Bastiaan Haange, Guylaine Miotello, Johan S. Sáenz, Maximilian Wolf, Felipe Bastida, Simon Devos, Guillermina Hernandez-Raquet, Jana Seifert, Paul Wilmes, Tim Van Den Bossche, Benoit J. Kunath, Robert Heyer, Nico Jehmlich, Dirk Benndorf, Robert L. Hettich, Jean Armengaud, Sergio Uzzau

**Affiliations:** Department of Biomedical Sciences, University of Sassari, Sassari, Italy; Unit of Microbiology and Virology, University Hospital of Sassari, Sassari, Italy; Multidimensional Omics Analyses Group, Leibniz-Institut für Analytische Wissenschaften – ISAS – e.V., Dortmund, Germany; Département Médicaments et Technologies pour la Santé (DMTS), SPI, Université Paris-Saclay, CEA, INRAE, Bagnols-sur-Cèze, France; Biosciences Division, Oak Ridge National Laboratory, Oak Ridge, TN, USA; Department of Molecular Toxicology, Helmholtz-Centre for Environmental Research – UFZ GmbH, Leipzig, Germany; Institute of Animal Science, University of Hohenheim, Stuttgart, Germany; HoLMiR—Hohenheim Center for Livestock Microbiome Research, University of Hohenheim, Stuttgart, Germany; Multidimensional Omics Analyses Group, Faculty of Technology, Bielefeld University, Bielefeld, Germany; Department of Soil and Water Conservation and Organic Waste Management, CEBAS-CSIC, Murcia, Spain; VIB-UGent Center for Medical Biotechnology, VIB, Ghent, Belgium; Department of Biomolecular Medicine, Faculty of Medicine and Health Sciences, Ghent University, Ghent, Belgium; VIB Proteomics Core, VIB, Ghent, Belgium; Toulouse Biotechnology Institute (TBI), Université de Toulouse, CNRS, INRAE, INSA, Toulouse, France; Luxembourg Centre for Systems Biomedicine, University of Luxembourg, Esch-sur-Alzette, Luxembourg; Department of Life Sciences and Medicine, University of Luxembourg, Esch-sur-Alzette, Luxembourg; Applied Biosciences and Process Engineering, Anhalt University of Applied Sciences, Köthen, Germany; Bioprocess Engineering, Otto von Guericke University, Magdeburg, Germany

**Keywords:** Metaproteomics, fecal microbiome, sample stabilization, mass spectrometry, CAMPI

## Abstract

**Background:** Fecal samples are widely used as a proxy for studying gut microbiome composition in both human and animal research. Fecal metaproteomics provides valuable insights by tracking changes in the relative abundance of microbial taxa and their protein functions. To ensure reliable results, it is crucial to minimize alterations in the metaproteome occurring from sample collection to protein extraction. Therefore, employing effective stabilization methods is essential to preserve the integrity of the fecal metaproteome from sample collection to laboratory analysis, particularly over long distances or when rapid freezing options are not readily available. In line with these needs, the second edition of the Critical Assessment of MetaProteome Investigation (CAMPI-2) was specifically focused on testing sample stabilization protocols to be applied before metaproteomic analysis.

**Results:** This collaborative multicenter study assessed the ability of five different stabilization methods, based on two commercial devices and three specific reagents (acetone, lithium dodecyl sulfate, and an RNAlater-like buffer), respectively, to stabilize the fecal metaproteome during room-temperature storage (14 days) and shipment to mass spectrometry facilities. The five methods were tested simultaneously by eight different laboratories across Europe, using aliquots from the same fecal sample. After protein extraction and digestion, duplicate aliquots of the resulting peptides were analyzed independently by two mass spectrometry facilities at distinct international locations. Analysis of the mass spectrometric data using two different search engines revealed that the fecal metaproteome profile differed considerably depending on the stabilization method used in terms of richness, alpha and beta diversity, reproducibility, and quantitative distribution of main taxa and functions. Although each method showed unique strengths and weaknesses, a commercial swab-based device stood out for its remarkable reproducibility and ranked highest for most of the metrics measured.

**Conclusions:** CAMPI-2 allowed a robust evaluation of five different methods for preserving fecal metaproteome samples. The present investigation provides useful data for the design of metaproteomics and multi-omics studies where fecal sampling cannot be immediately followed by long-term storage at −80°C. Further optimization of the tested protocols is necessary to improve stabilization efficiency and control bias in the taxonomic and functional profile of the gut microbiome.

## Background

Metaproteomics is a powerful technology that enables functional and taxonomic analysis of microbial communities through liquid chromatography coupled to high-resolution tandem mass spectrometry (LC-MS/MS) [1]. Notably, metaproteomics is highly complementary to other meta-omics approaches, as it provides an additional layer of information by linking the functional capacity of the microbial communities to their actual phenotype. To enhance this technology, the metaproteomics research community has initiated a community-driven, multi-laboratory effort to compare workflows, under the umbrella of the Metaproteomics Initiative [2]. The first edition of the Critical Assessment of MetaProteome Investigation (CAMPI) examined the effects of sample preparation protocols, MS parameters, and bioinformatics approaches, demonstrating similar functional profiles across workflows [3]. The second edition of the Critical Assessment of MetaProteome Investigation, CAMPI-2, focuses on testing different sample stabilization protocols to be applied before metaproteomic analysis.

Fecal samples are easily accessible and self-collectable specimens that are commonly used as a proxy for investigating gut microbiome composition in human and animal studies [4]. Yet, dealing with highly complex samples such as human feces is particularly challenging, as environmental stimuli strongly influence the target of investigation. Therefore, nucleic acids and/or proteins should be extracted from fresh feces as soon as possible to avoid bias due to microbial overgrowth [5]. However, performing extraction immediately after sample collection is often impossible. Temperature, humidity, pH, and oxygen concentration are known to have a relevant and rapid impact on microbial gene expression and protein synthesis/degradation, which can be affected after a short exposure to a new environment [6]. Thus, the choice of storage conditions from the time of sampling to extraction is of paramount importance when dealing with metaproteomic analyses. Flash freezing of the fecal sample in liquid nitrogen is not a practical procedure, and in many sampling situations it is not even feasible. Hence, freezing the fecal sample at a minimum of −80 °C shortly after collection is currently considered the “gold standard” in metaproteomics [7]. However, the length and conditions of the period between sampling and freezing can vary considerably, especially in the case of population studies based on self-collection, adding a source of technical variability. In addition, storing and shipping frozen fecal samples can also be logistically challenging, especially in studies involving individuals and cohorts without access to temperature-controlled storage and transport systems. All these issues underscore the need for straightforward and robust sample stabilization protocols that allow for the exchange of fecal samples at room temperature between collection sites and laboratories, even over long distances [5]. Therefore, the choice of sample collection protocols and storage conditions is a key requirement for application in population-based fecal metaproteomic studies, and their reproducibility is essential to ensure data quality.

Fecal sample stabilization methods suitable for metaproteomic studies are expected to meet at least four key requirements: i) efficiency, i.e. the ability to stop as soon as possible any potential metaproteome profile shift during the environmental transition from the rectal ampulla to the fecal sample container; ii) stability, i.e. the ability to stabilize the collected sample at room temperature until it is shipped to the main laboratory, where it is frozen at −80°C while the entire set of samples is being collected; iii) ease of use, allowing untrained individuals to perform self-sampling of feces while maintaining high levels of reproducibility; iv) lack of interference with downstream procedures, including protein digestion, chromatographic separation of peptides, and MS analyses [7]. Routine protocols should also be affordable and rapid. Since multi-omics designs are known to provide a deeper understanding of the biological system under study [8], an ideal stabilization method should also be compatible with the analysis of other classes of molecules (i.e. DNA, RNA, metabolites).

Among the already validated methods for storing cells and tissues while protecting the stability of nucleic acids, RNAlater has been tested for handling human fecal samples in metaproteomic studies with conflicting results [9]. In addition, pre-analytical sample stabilization using various swab-based devices has become common in recent years, allowing for more robust and standardized protocols for collection, room-temperature storage, and transport to the molecular analysis laboratory of clinical specimens, including feces. Thorough inactivation of biological activity allows fecal samples to be conveniently stored, transported, and disposed of as non-hazardous waste without requiring special handling. While these swab-based methods have been tested for use in metagenomic studies [10], also in comparison to other methods [11], to the best of our knowledge their suitability for fecal metaproteome analyses has not yet been systematically examined. In addition, little is known about potential biases (e.g. proteolysis or expression of novel proteins) that this type of sample treatment might introduce in human gut metaproteome profiling. Furthermore, stabilization of fecal samples can also be performed through methods commonly employed for rapid protein extraction from microbial communities by denaturation/precipitation, including protocols based on lithium dodecyl sulfate (LDS), sodium dodecyl sulfate (SDS), and acetone [12–14].

Great care should be taken in choosing methods for the stabilization and storage of fecal specimens and for their experimental assessment. Critical comparison of stabilized metaproteome samples must be performed under real-world conditions, by evaluating their stability over time at room temperature and assessing their quality when subjected to MS analysis. Evaluation of protocol efficiency and robustness also requires the participation of a significant number of laboratories to control inter-operator variability in all pre-analytical steps. Here, we tested five different stabilization methods, based on reagents or devices already available to the proteomics and/or the microbiome research communities, in an inter-laboratory trial. Specifically, we evaluated the ability of these methods to preserve the fecal metaproteome profile after 14 days of room-temperature storage, followed by shipment of stabilized samples to MS facilities without delivery priority (5-7 business days) and without control of transport temperatures.

## Results

### Experimental design

The experimental design of the study is depicted in **Fig. 1**. In the “Aliquoting lab” (aLab; University of Sassari, Italy), a fecal sample provided by a healthy volunteer was collected and divided into aliquots, which were immediately stored at −80°C. The fecal aliquots were then shipped on dry ice to eight different “Stabilization labs” (sLabs) across Europe.

**Fig. 1.**
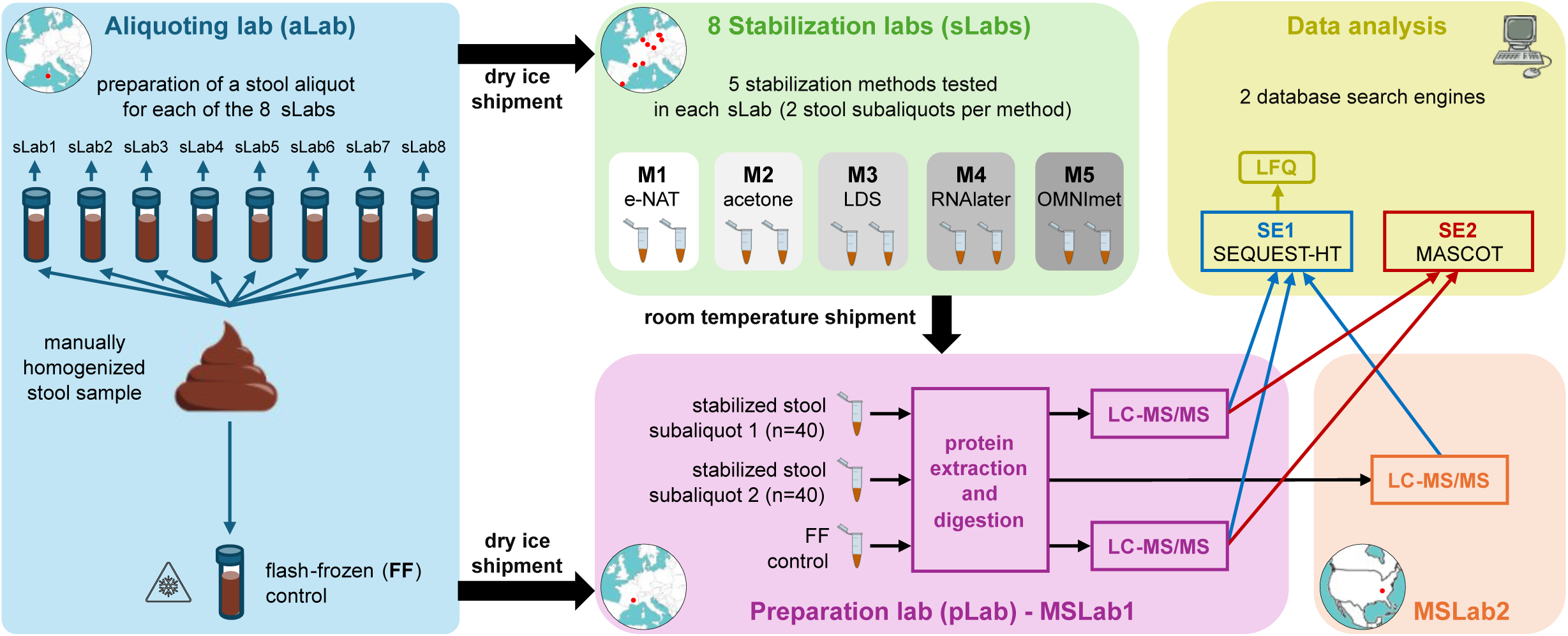
Experimental design of the study.

Each sLab processed ten fecal aliquots using five different stabilization methods (two fecal aliquots per method), based on the e-NAT^®^ collection and preservation system (M1), acetone (M2), the LDS sample buffer (M3), an RNAlater-like buffer (M4), and the OMNImet^®^•GUT device (M5), respectively. The stabilized samples were kept at room temperature for 14 days and then shipped to the “Preparation lab” (pLab; CEA, France). In parallel, a fecal aliquot kept frozen at −80°C was shipped on dry ice from the aLab to the pLab and considered as the flash-frozen (FF) control.

In the pLab, stabilized samples and the FF control were subjected to protein extraction and digestion. Unfortunately, technical issues were encountered during the processing of M5 samples, with protein precipitation failing to yield a pellet, probably due to the high ethanol content of the stabilizing medium. As a result, protein digestion could only be successfully performed on six M5 samples.

The peptide mixtures obtained from the first subaliquot of all the stabilized samples (one replicate per each sLab-method combination) and from the FF controls (three run replicates) were analyzed by LC-MS/MS in the same lab (MSLab1; ProGénoMix platform, CEA, France), while those obtained from the second subaliquot of the stabilized samples were sent to a second MS lab (MSLab2; ORNL, USA) for parallel LC-MS/MS analysis. Due to the problems described above, three of the M5 samples could only be analyzed in the MSLab1 because of the low amount of peptides obtained during sample preparation.

The MS files produced by MSLab1 and MSLab2 were searched against a combination of microbial and human sequence databases using SEQUEST-HT as the search engine (SE1), resulting in two datasets named MSLab1-SE1 and MSLab2-SE1, respectively. Label-free quantification (LFQ) data were retrieved from both SE1-based datasets. In addition, to ensure that the results were not dependent on a specific data analysis pipeline, the MSLab1 files were also analyzed using MASCOT as the search engine (SE2) and a different approach to control the false-discovery rate (FDR), producing the MSLab1-SE2 dataset.

### Identification metrics according to different LC-MS/MS and database search pipelines

We first measured the global identification yield achieved with the five methods by calculating the number of peptide-to-spectrum matches (PSMs; **Fig. 2**, left) and of unique peptide sequences (**Fig. 2**, middle) identified after searching against the microbial/human databases. The FF control always achieved the highest number of identifications. All three datasets show that M1 performed best among stabilized samples, with an average of −27% PSMs and −15% peptides compared to FF. M3 came in second, followed by M2, M4, and M5. In quantitative terms, when compared to M1 (average between datasets), M3 provided 20% fewer PSMs and 22% fewer peptides; M2 provided 59% fewer PSMs and 72% fewer peptides; M4 provided 75% fewer PSMs and 76% fewer peptides; and M5 provided 98% fewer PSMs and 98% fewer peptides. It is worth noting that considerable differences between the stabilization methods were also observed with respect to the mean number of MS/MS spectra acquired (**Fig. S1**), indicating that specific moieties might have interfered with MS identification and/or peptide quantitation prior to LC-MS/MS analysis.

**Fig. 2.**
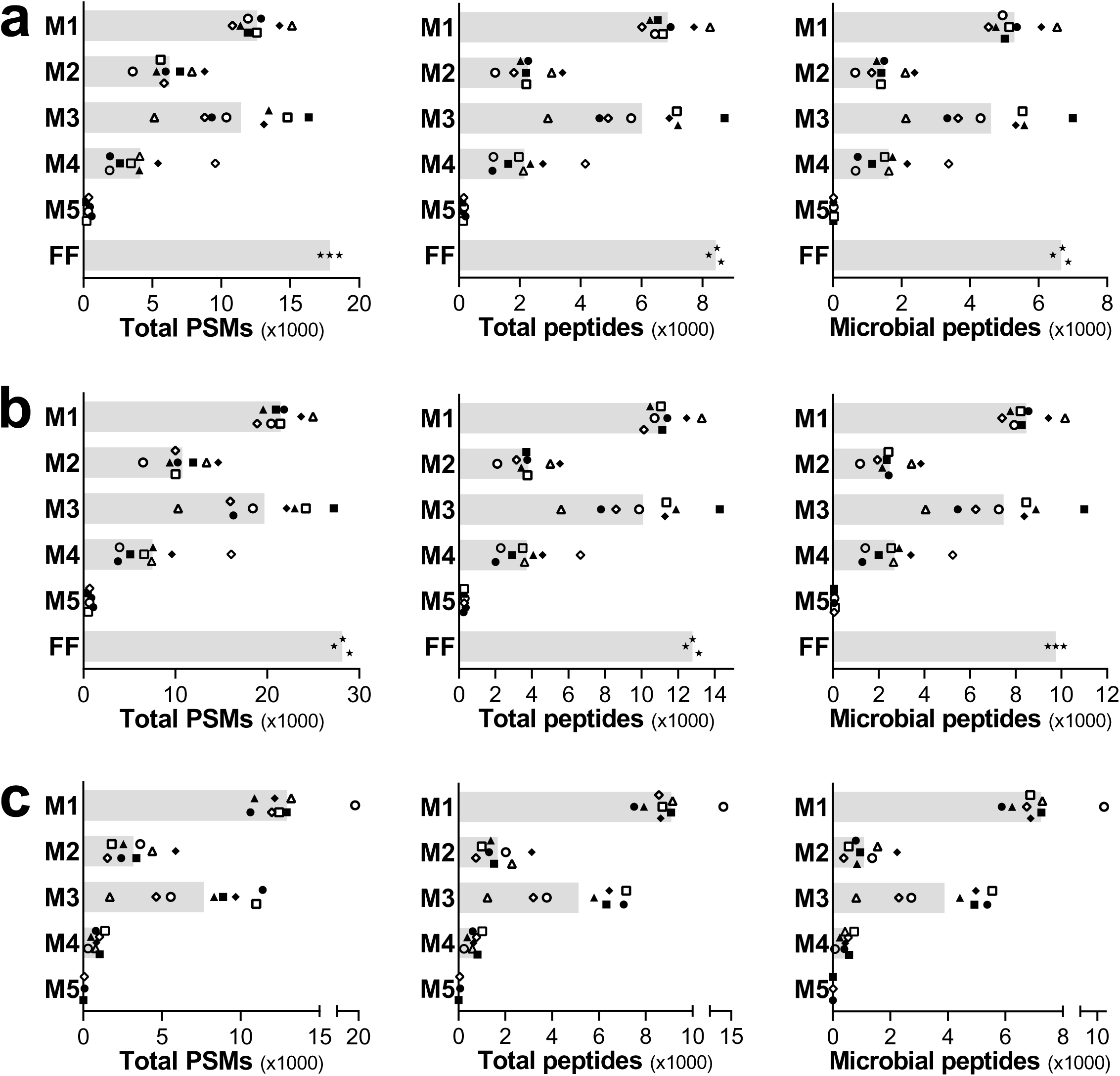
Identification metrics according to the MSLab1-SE1. (**a**), MSLab1-SE2 (**b**), and MSLab2-SE1 (**c**) datasets. Scatter bar plots illustrate the total number of PSMs identified (left), the total number of distinct peptide sequences identified (middle), and the number of distinct microbial peptide sequences identified (right) in the same fecal sample after processing with the five stabilization methods being compared (M1-M5) or after flash freezing (FF). Spectra were searched against a combination of microbial and human sequence databases; among the identified peptides, those taxonomically assigned to bacteria, archaea, or viruses were considered as microbial, and their numbers are shown in the right plots. Each shape marks a different sLab replicate, while the grey bars indicate the mean value for each method.

We then focused our analysis on the microbial peptide sequences, i.e. those unambiguously assigned to the superkingdoms Bacteria, Archaea, or Viruses according to the Unipept taxonomic annotation [15] (**Fig. 2**, right). The trends observed for microbial peptides were globally similar to those described above for the combination of microbial and human peptides. Since the number of microbial peptides identified in the samples stabilized with M5 was extremely low and clearly incomparable to the other methods (as well as 200 times lower than the FF control) according to all three datasets, M5 was excluded from the subsequent analyses.

Furthermore, UpSet plots generated after merging the microbial identifications obtained with all replicates for each group revealed that a large number of peptides were unique to FF, M1, and M3, in descending order, whereas most peptides in M2 overlapped with those identified by the other methods (**Fig. S2**). Moreover, a considerable fraction of peptides was shared between M1 and M3.

In addition, we investigated the degree of identification overlap between each stabilization method and the FF control, according to the MSLab1-SE1 and MSLab1-SE2 datasets (**Fig. 3**). When evaluating the intersection between the sLab replicates of each stabilization method and FF in terms of microbial peptides identified, M1 and M3 were observed to have the highest number of peptides in common with FF, followed by M2 and M4. However, over 50% (MSLab1-SE1) and 60% (MSLab1-SE2) of the peptides identified with M1, M3, and M4 were not identified at all in the FF control, as compared to only 13% (MSLab1-SE1) and 28% (MSLab1-SE2) of M2 identifications. To verify whether the peptides detected in the stabilized fecal samples but not in the FF control were mainly of low abundance, we recalculated the intersection between the stabilization methods and the FF control after setting different thresholds based on the minimum number of PSMs detected per peptide (namely, 3, 5, or 10). For both datasets, as the PSM-based threshold increased, the mean percentage of peptides detected in the stabilized samples but not detected in FF decreased, particularly for M1 and M3. This suggests that most medium-to high-abundance peptides identified in these stabilized samples were also present in the FF control. Notably, M2 displayed the highest mean number of microbial peptides detected with more than 10 PSMs, both in absolute terms and in those shared with FF.

**Fig. 3.**
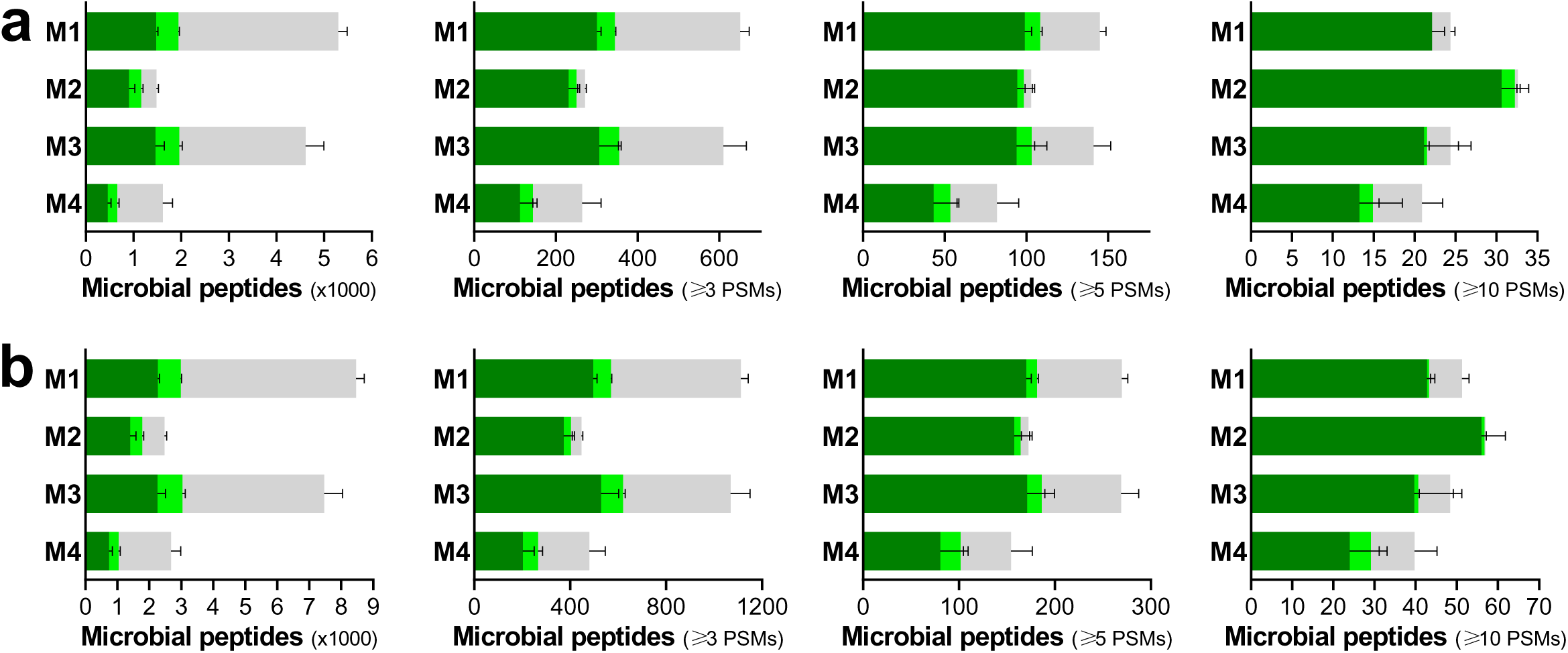
Overlap of microbial peptide identifications between each of the four stabilization methods and the FF control, according to the MSLab1-SE1. (**a**) and MSLab1-SE2 (**b**) datasets. Each bar indicates the mean number of peptides identified in the eight sLab replicates, with error bars indicating the standard error of the mean. Bars are divided into three areas based on the degree of overlap with the FF control: the dark green areas include peptides that were also identified in all three FF replicates, the light green areas include peptides that were also identified in 1 or 2 FF replicates, and the gray areas include peptides that were not identified in any FF replicate. The left plots refer to all the peptides identified, while the others were generated after setting increasing thresholds based on the minimum number of PSMs detected per peptide (namely 3, 5, and 10).

### Diversity and reproducibility of metaproteome profiles

After verifying that the main identification metrics did not appear to be substantially biased by the specific MS platform or search engine used, the MSLab1-SE1 dataset was selected as the sole dataset for all subsequent analyses. LFQ abundances measured for microbial peptides were then used to compute several quantitative metrics, including alpha and beta diversity and method reproducibility.

First, we calculated Shannon’s index to evaluate the level of alpha diversity reached with the different stabilization methods. As shown in **Fig. 4a**, similar alpha-diversity values were measured for M1, M3 and FF, followed by M4 and M2. Interestingly, M1 also showed very low variability between sLab replicates.

**Fig. 4.**
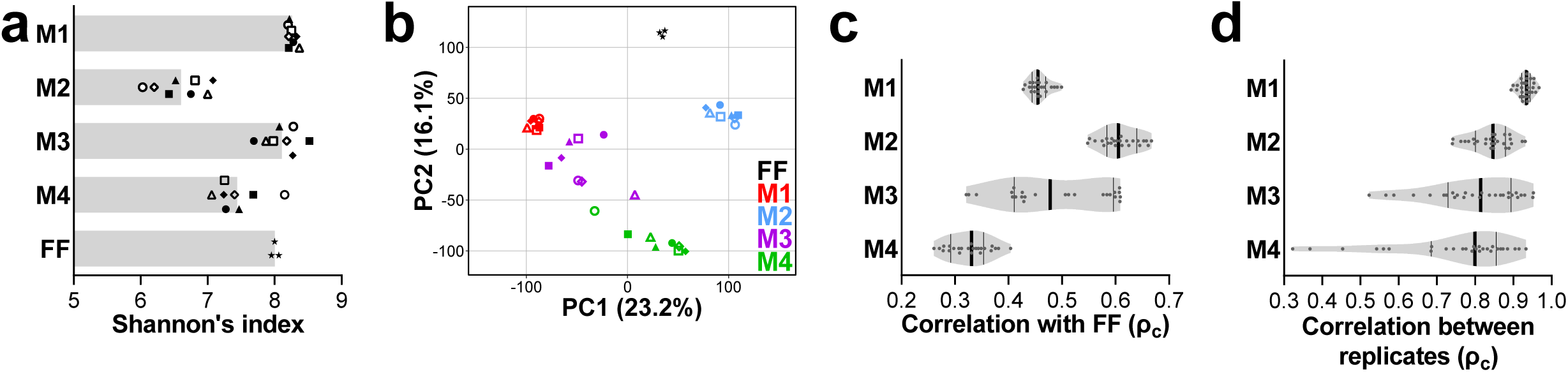
Diversity and correlation analyses based on the distribution of peptide LFQ abundance values measured in the same fecal sample after processing with four stabilization methods (M1-M4) or after flash freezing (FF), according to the MSLab1-SE1 dataset. **a** Scatter bar plot illustrating the peptide alpha-diversity values measured according to the Shannon’s index. Each shape marks a different sLab replicate, while the grey bars indicate the mean value for each method. **b** Principal component analysis (PCA) plot. Each shape marks a different sLab replicate, while each method is marked with a different color. The percentages of variation explained by the first two components are shown in the x– and y-axes, respectively. **c** Violin plot showing the distribution of the concordance correlation coefficient (ρ_c_) calculated between the metaproteome profiles obtained with each of the stabilization methods under comparison (M1-M4) and that of the flash-frozen (FF) control. Each dot corresponds to the correlation value calculated between an FF replicate and a method (sLab) replicate. The vertical thick black line indicates the median of the distribution, while thinner lines indicate the upper and lower quartiles. **d** Violin plot showing the correlation between sample replicates based on the concordance correlation coefficient (ρ_c_), as a measure of method reproducibility. Each dot corresponds to the correlation between two sLab replicates of a specific method. The vertical thick black line indicates the median of the distribution, while thinner lines indicate the upper and lower quartiles.

To investigate beta diversity between samples, principal component analysis (PCA) was performed. The PCA plot of **Fig. 4b** confirmed the much higher reproducibility of M1 compared to the other stabilization methods (especially M3), along with a globally lower distance of FF with the M2 cluster compared to the M1 and M3 clusters. Furthermore, the variable ‘method’ was found to have a much stronger impact on the metaproteome profile than the variable ‘sLab’.

Furthermore, we correlated the peptide LFQ abundance values between the 8 sLab replicates of each stabilization method and the run replicates of the FF control, as a further measure of quantitative comparability between their metaproteome profiles. As shown in **Fig. 4c**, the highest median concordance correlation coefficient (ρ_c_) value was observed for the M2 *vs* FF comparison (0.61), followed by M3 (0.49), M1 (0.46), and M4 (0.33).

As a more specific measure of technical reproducibility, we correlated the peptide LFQ abundance values obtained in the different sLab replicates with each other, separately for each stabilization method (**Fig. 4d**). M1 had by far the highest concordance correlation coefficient value (mean ρ_c_ 0.93), thus proving to be the most reproducible method, followed by M2 (0.84), M3 (0.80), and M4 (0.74).

Finally, to estimate the amount of hydrophobic proteins preserved with the different methods, the relative abundance of peptides with a GRAVY score higher than 0.5, which is considered as a reasonable threshold for hydrophobicity [16,17], was calculated (**Fig. S3**). M3 emerged as the method providing the highest amount of hydrophobic peptides (with around 30% increase compared to FF), while M4 values were those closest to the FF control.

### Richness and abundance distribution of taxonomic profiles

To go deeper on the taxonomic composition of the metaproteome profiles obtained with the different stabilization methods and in the FF control, we first evaluated the peptide annotation yield at different taxonomic levels (**Fig. S4**). A slight decrease in the taxonomic annotation rate of the identified peptides was observed at all hierarchical levels in the samples subjected to the stabilization methods compared to FF, with the average decrease ranging from 10% for M3 to 5% for M2. Taking the genus level as an example, the percentage of peptides with a genus-specific annotation ranged from 37.5% for FF down to 27.7% for M3.

Then, we examined peptide identification data to infer the number of distinct taxa (i.e. phyla, classes, orders, families, genera, and species) detected with each of the different methods, as a measure of taxonomic richness. As shown in **Fig. 5a**, a higher taxonomic richness was achieved on average in samples stabilized with M3 compared to M1 and FF (which were similar to each other), while lower levels were measured for M2 and M4, in line with the trends seen for the peptide identification yield. As shown in the UpSet plots of **Fig. 5b**, 100 taxa were identified with all methods, whereas 59 and 53 taxa were unique to M3 and M1, respectively. Further 35 taxa were shared between M1 and M3 and not detected in the other sample groups. The intersections between each stabilization method and the FF control showed that M3 and M1 provided the highest number of taxa shared with FF (120 and 115, respectively), as illustrated in **Fig. 5c**. However, another 67 and 56 of the taxa identified with M3 and M1, respectively, were not detected in FF. Similar to what described above for peptides, we set different thresholds based on the minimum number of PSMs detected per taxa to investigate how this impacted on the degree of overlap between the stabilization methods and the FF control in terms of taxonomic richness. As a result, the percentage of taxa not shared with FF decreased drastically as the PSM-based threshold increased. This was particularly evident for M1, where the percentage of taxa not in common with the FF control decreased to 6% and 0% when considering the microbial taxa detected with at least 3 and 5 PSMs, respectively. Interestingly, *Dialister* was detected in most of the M1 and M3 replicates (with an average of 4 PSMs in the latter group) but was not identified in any of the FF replicates. A similar behavior was found for *Eisenbergiella* with respect to M2 and M4. On the other hand, among the microbial genera found in all FF replicates with at least 3 PSMs on average, *Romboutsia* was not detected in M1, *Faecalibacillus* was not detected in either M1 or M2, *Odoribacter* was not detected in either M1 or M4, while *Vescimonas* and *Akkermansia* were not detected in M4. The overlap between the four stabilization methods and the FF control distinctly calculated at the family, genus, and species levels is shown in **Fig. S5**.

**Fig. 5.**
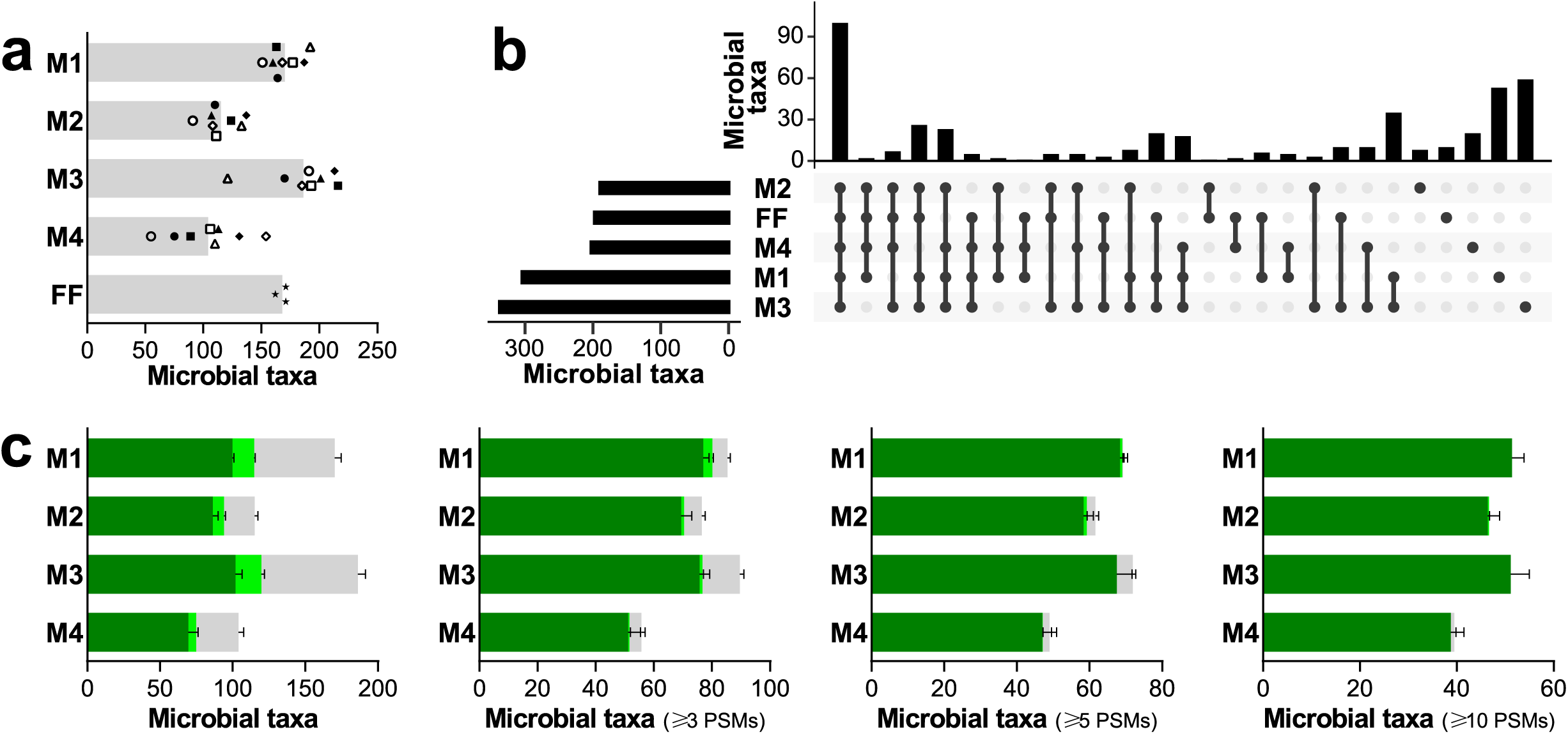
Taxonomic richness measured in the same fecal sample after processing with four stabilization methods (M1-M4) or after flash freezing (FF), according to the MSLab1-SE1 dataset. Data were based on the number of distinct microbial taxa identified across the six main hierarchical levels (from phyla down to species). **a** Scatter bar plot illustrating the number of taxa identified as a measure of taxonomic richness. Each shape marks a different sLab replicate, while the grey bars indicate the mean value for each method. **b** UpSet plot showing the identification overlap between the different methods in terms of microbial taxa. Taxa identified in the different replicates were merged for each method. Horizontal bars indicate the set size, i.e. the total number of taxa identified for each method (sorted by increasing set size from top to bottom). Vertical bars illustrate the intersection size, i.e. the number of taxa shared between the different methods (sorted by decreasing degree of overlap from left to right), according to the black dot(s) below. **c** Overlap of the identified taxa between the four stabilization methods and the FF control. Each bar indicates the mean number of taxa identified between sLab replicates, with error bars indicating the standard error of the mean. Bars are divided into three areas based on the degree of overlap with the FF control: the dark green areas include taxa that were also identified in all three FF replicates, the light green areas include taxa that were also identified in 1 or 2 FF replicates, and the gray areas include taxa that were not identified in any FF replicate. The left plot refers to all the taxa identified, while the others were generated after setting increasing thresholds based on the minimum number of PSMs detected per taxa (namely 3, 5, and 10).

Peptide LFQ values were then aggregated to calculate the relative abundances of the main taxa present in the different sample groups. **Fig. 6** shows the relative abundance of the top 5 microbial phyla (**a**), the top 15 microbial families (**b**), and the top 20 microbial genera (**c**) measured in the same fecal sample after processing with the four stabilization methods and after flash freezing, selected based on their mean relative abundance in the FF group. As general considerations based on these data, M1 exhibited the highest degree of similarity to the FF taxonomic profile, M2 and M3 profiles varied considerably between replicates, while dramatic changes relative to FF were observed for M4. Each stabilization method showed specific taxonomy distribution biases compared to the FF control. Bacteroidota, Bacteroidaceae and their genera *Bacteroides* and *Phocaeicola* were all relatively less abundant in M4. Within Bacteroidota, Rikenellaceae and its genus *Alistipes* were relatively decreased in all methods (especially M4). On the other hand, Bacillota were relatively more abundant in M4. Among the Bacillota families, Coprobacillaceae (and its genus *Faecalibacillus*) were relatively decreased in all methods, Peptostreptococcaceae were relatively less abundant in M1 and M3, and Streptococcaceae were relatively increased in M1 and M3. Among the Bacillota genera, *Anaerotruncus* and *Evtepia* were relatively less abundant in all methods, while *Clostridium* was relatively increased in all methods; *Dorea* was relatively more abundant in M4 and decreased in all other methods. Among the less represented phyla, Thermodesulfobacteriota (and its family Desulfovibrionaceae) were relatively increased in M2, while Pseudomonadota were relatively increased in M2 and M4.

**Fig. 6.**
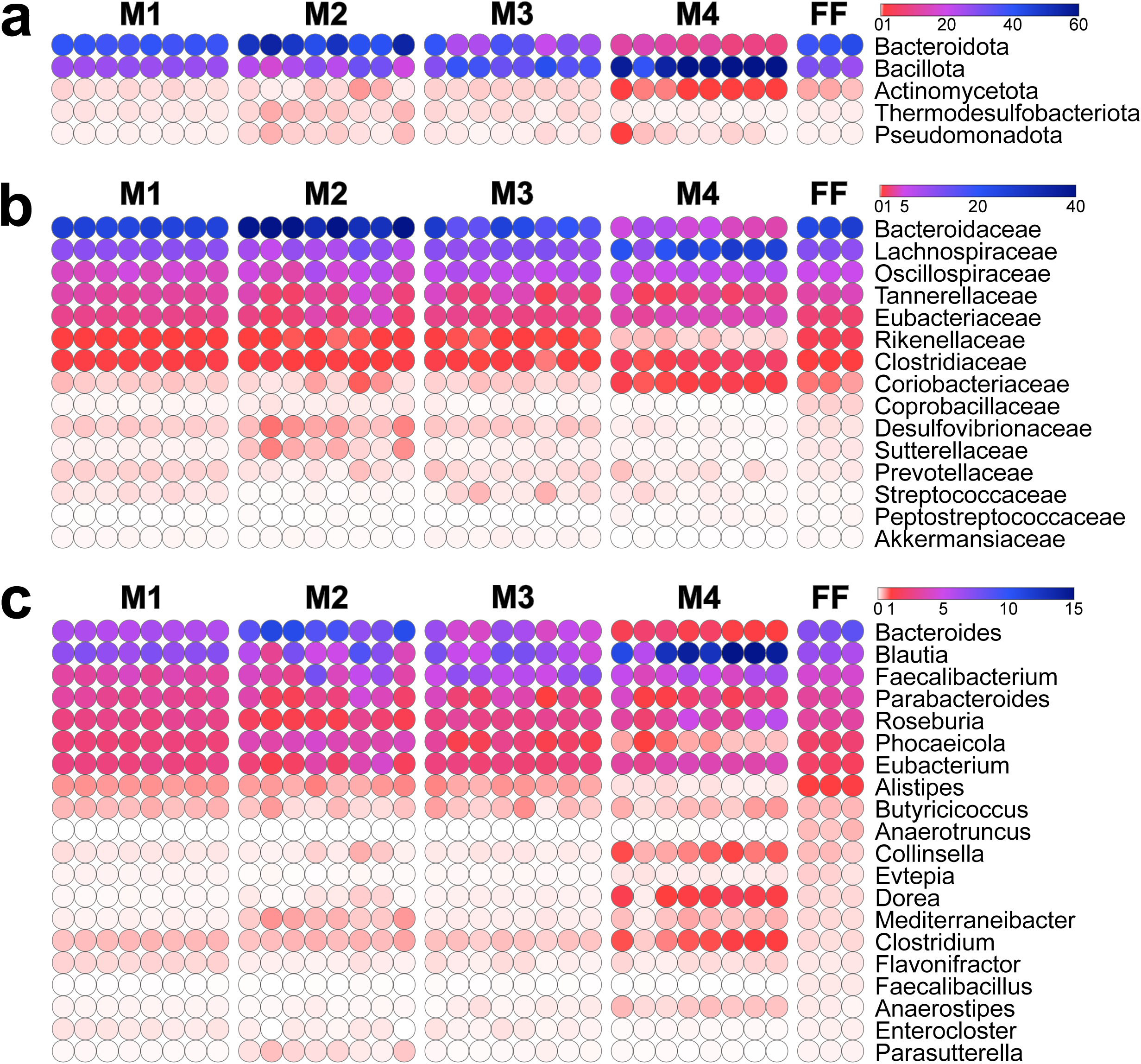
Relative abundances of the top 5 microbial phyla. (**a**), the top 15 microbial families (**b**), and the top 20 microbial genera (**c**), selected based on their mean relative abundance in the FF group, according to the MSLab1-SE1 dataset. Abundances were calculated by summing LFQ values of all peptides unambiguously assigned to each taxon and are expressed as a color gradient according to the legend in the upper right corner of each heatmap. Each dot represents a different sample replicate.

Moreover, phylum-level data were used to calculate the abundance ratio between Bacillota and Bacteroidota (formerly known as the Firmicutes/Bacteroidetes ratio), regarded as a relevant metric able to summarize microbiome compositional changes related to diet, age, and disease [18–20]. According to the log ratio values shown in **Fig. S5**, M1 provided the value closest to the FF control (−0.14 and –0.16, respectively). Increasingly distant values were measured for M2 (−0.30 on average), M3 (0.09 on average) and M4 (0.75 on average).

### Richness and abundance distribution of functional profiles

Next, we analyzed the functional composition of the microbial metaproteomes obtained using the stabilization methods and the FF control. Firstly, we examined the functional annotation yield. As shown in **Fig. S6**, the average percentage of microbial peptides successfully assigned to a specific KEGG orthology (KO) function was notably higher for M4, M3, and M1 (in descending order) compared to FF.

Then, peptide identification data were parsed to calculate the level of functional richness achieved with the different stabilization methods, in terms of the number of distinct microbial KO functions identified (**Fig. 7a**). Functional richness levels were comparable between M1 and M3, in both cases slightly higher than the FF control and considerably higher than the other stabilization methods. Furthermore, as shown in the UpSet plot of **Fig. 7b**, the largest intersection between methods was the one containing the functions unique to M1 and M3, followed by the one containing the functions detected in all sample groups. Regarding the overlap between each stabilization method and the FF control (**Fig. 7c**), the highest number of functions shared with FF was observed in the samples stabilized with M1 and M3. However, approximately 45% of the functions identified with these two methods were not detected in the control. On the contrary, 93% of the microbial functions detected in the samples stabilized with M2 were in common with FF. After setting increasing thresholds based on the minimum number of PSMs detected per function, we did not observe any clear reduction in the percentage of peptides shared with the FF control, indicating that a considerable fraction of medium/high-abundance microbial functions were identified only in the stabilized fecal aliquots, particularly in M1 and M3 samples. Specifically, 58 and 43 functions identified with more than 10 PSMs on average in M1 and M3 samples, respectively, of which 35 shared between the two sample groups, were not identified in FF. These included DnaK and other molecular chaperones, RNA polymerase subunits, numerous ribosomal proteins, and several proteins involved in carbohydrate transport and metabolism and in oxidative phosphorylation. On the other hand, 54 functions detected with more than 10 PSMs on average in FF samples were not identified in one or more stabilized sample groups (of which 47, 17, 9, and 3 were undetected in M4, M2, M1, and M3, respectively). These included several enzymes involved in pyruvate, purine, glycan, and amino acid metabolism.

**Fig. 7.**
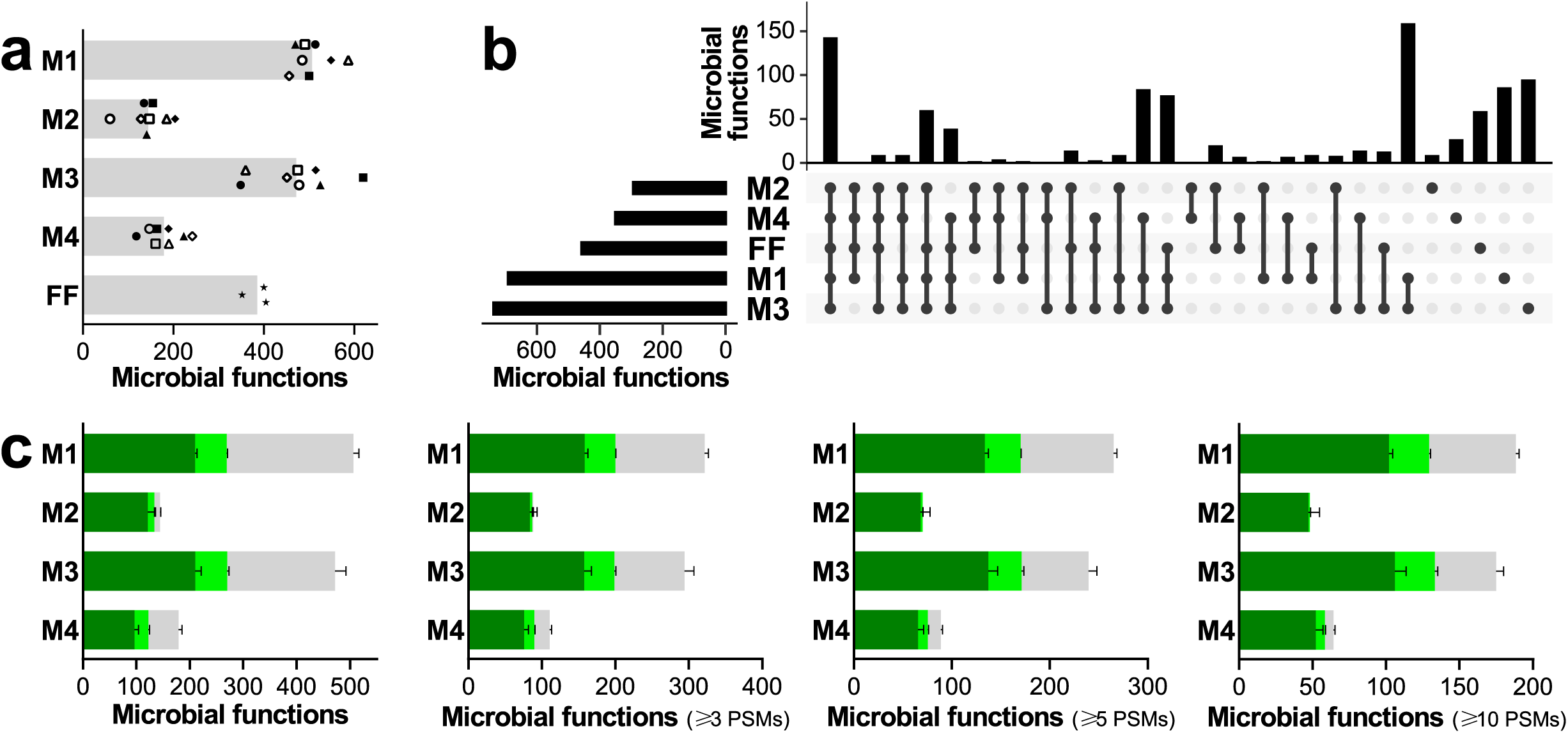
Functional richness measured in the same fecal sample after processing with four stabilization methods (M1-M4) or after flash freezing (FF), according to the MSLab1-SE1 dataset. Richness was intended as the number of distinct microbial KEGG KO functions identified. **a** Scatter bar plot illustrating the number of functions identified as a measure of functional richness. Each shape marks a different sLab replicate, while the grey bars indicate the mean value for each method. **b** UpSet plot showing the identification overlap between the different methods in terms of microbial functions. Functions identified in the different replicates were merged for each method. Horizontal bars indicate the set size, i.e. the total number of functions identified for each method (sorted by increasing set size from top to bottom). Vertical bars illustrate the intersection size, i.e. the number of functions shared between the different methods (sorted by decreasing degree of overlap from left to right), according to the black dot(s) below. **c** Overlap of the identified functions between the four stabilization methods and the FF control. Each bar indicates the mean number of functions identified between sLab replicates, with error bars indicating the standard error of the mean. Bars are divided into three areas based on the degree of overlap with the FF control: the dark green areas include functions that were also identified in all three FF replicates, the light green areas include functions that were also identified in 1 or 2 FF replicates, and the gray areas include functions that were not identified in any FF replicate. The left plot refers to all the functions identified, while the others were generated after setting increasing thresholds based on the minimum number of PSMs detected per function (namely 3, 5, and 10).

As a further measure of the functional richness, we calculated the number of KO functions detected with the different methods and mapping to one of the main KEGG metabolic pathways, namely the one named “Carbon metabolism” (map01200). As illustrated in **Figs. S7-S11**, similar results were obtained for M3 and M1 (74 and 71 mapped enzymes, respectively), followed by FF and M4 (50 and 48 mapped enzymes, respectively).

Finally, we went deeper into the investigation of the functional profile of the metaproteomes obtained with the different stabilization methods by aggregating the LFQ abundance values of microbial peptides according to their Cluster of Orthologous Group (COG) and KO annotations. **Fig. 8a** illustrates the relative abundance distribution of the 20 main COG functional categories, ordered based on their mean relative abundance in the FF group. Proteins involved in transport and metabolism of inorganic ions were relatively less abundant in the samples treated with all stabilization methods (especially M4) compared to the FF control, while those involved in translation, post-translational modification, and protein turnover were relatively more abundant in M1 and M3. A relative increase in proteins involved in cell motility and in coenzyme transport and metabolism was also observed in M2– and M4-treated samples, respectively. Interestingly, among peptides associated with a “ Cell motility” COG annotation, nearly all of those with the highest abundance values in M2 lack a KO annotation, likely corresponding to uncharacterized/unknown proteins. Furthermore, M1 exhibited a relative decrease in proteins involved in transport and metabolism of carbohydrates.

**Fig. 8.**
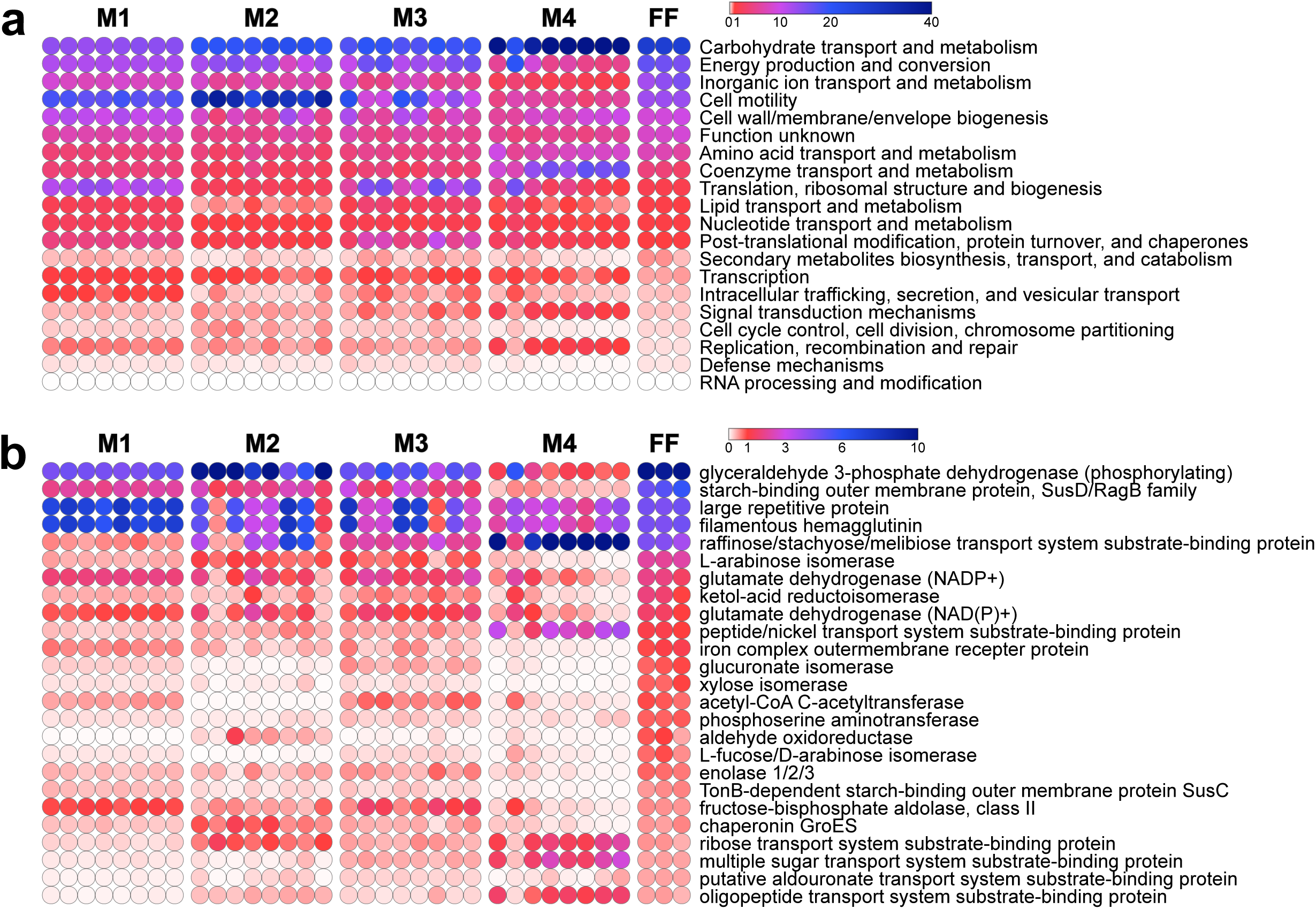
Relative abundances of the top 20 microbial Cluster of Orthologous Groups (COGs) (**a**) and of the top 25 microbial KO functions (**b**), selected based on their mean relative abundance in the FF group, according to the MSLab1-SE1 dataset. Abundance values are expressed as a color gradient according to the legend in the upper right corner of each heatmap. The abundance of each COG (**a**) or KO (**b**) function was calculated as the sum of the LFQ abundances of all peptides functionally assigned to that COG/KO. Each dot represents a different sample replicate.

The abundance profiles of the top 25 KO functions, selected based on their mean relative abundance in the FF samples, are shown in **Fig. 8b**. Several enzymes with high relative abundance in the FF control were relatively decreased in the samples treated with the stabilization methods, mostly sugar isomerases. Among specific functions relatively increased in samples treated with specific stabilization methods (**Supplementary Data 1**), we can mention flagellin for M1, several ribosomal subunits, chaperonins and elongation factors, as well as some short-chain fatty acid biosynthetic enzymes, for M1 and M3, transketolase for M2, and some transport system substrate-binding proteins for M4.

Although this study was mainly focused on the microbial fraction of the metaproteome, we also investigated the relative abundance distribution of the top 25 human proteins, selected based on their mean relative abundance in the FF samples (**Fig. S12**). Again, each stabilization method exhibited its specific increase and decrease trends with respect to the FF control.

## Discussion

In the earliest CAMPI (hereafter CAMPI-1), launched by the Metaproteomic Initiative [2], a multi-laboratory network of metaproteome researchers analyzed and compared the performances of a large variety of experimental procedures and computational pipelines to profile a human fecal sample and a simplified human gut microbiota model (SIHUMIx) [3]. CAMPI-1 results highlighted the overall similarity of the profiles obtained by different workflows of sample extraction, mass spectrometry, and bioinformatics analysis, with minor differences mainly in the levels of analytical depth due to the extent of sample fractionation and the characteristics of the mass spectrometers.

Notably, the structure and functions of a microbial community are governed by the interaction with its physical and biological environment, which can be easily altered well before the analytical steps of the metaproteomics workflows take place. Accordingly, CAMPI-2 was designed to address the importance of selecting appropriate sampling and stabilization protocols suitable for microbiome samples, including feces. Samples to be subjected to metaproteomic analysis are typically flash-frozen and stored at −80°C soon after collection [21]. Long-term storage of extracted proteins is not recommended since their stability is compromised compared to proteins in their intact native matrix [22]. To facilitate the short-term storage of fecal microbiome samples, especially in studies involving sample self-collection in participants’ homes, several methods have been evaluated in sequencing-based fecal microbiome investigations [10,23,24]. Among the most recognized stabilization methods for metagenomic analysis, the use of the RNAlater preservative has also been tested for its efficiency in stabilizing different types of metaproteomes [9,25,26]. The two studies that evaluated RNAlater for the stabilization of fecal samples prior to metaproteomic analyses have reported conflicting results. One study concluded that preservation of mouse feces in RNAlater at room temperature was as efficient as freezing [25], whereas the other study showed that storage in RNAlater at 4°C for 6 hours had a strong impact on human metaproteomic and metagenomic profiles [9]. Therefore, we decided to compare the reliability and robustness of RNAlater with four other methods based on different devices and reagents that had yet to be thoroughly validated for metaproteomic studies.

In this study, we report that sampling and stabilization procedures have a strong impact on the metaproteomic profiles. On the contrary, as expected from CAMPI-1 data, different mass spectrometers or search engines applied to the same method led to quite similar trends. Each of the different methods tested in this comparative study had its own strengths and weaknesses, as summarized in **Table 1**.

**Table 1.** Ranking of each stabilization method according to the main metrics measured in this study.

| Metric | Rank |  |  |  |  |
| --- | --- | --- | --- | --- | --- |
|  | M1 | M2 | M3 | M4 | FF |
| PSMs identified (MSLab1-SE1) | 1 | 3 | 2 | 4 | >1 |
| PSMs identified (MSLab2-SE1) | 1 | 3 | 2 | 4 |  |
| PSMs identified (MSLab1-SE2) | 1 | 3 | 2 | 4 | >1 |
| Peptide sequences identified (MSLab1-SE1) | 1 | 3 | 2 | 3 | >1 |
| Peptide sequences identified (MSLab2-SE1) | 1 | 3 | 2 | 4 |  |
| Peptide sequences identified (MSLab1-SE2) | 1 | 3 | 2 | 3 | >1 |
| Microbial peptides identified (MSLab1-SE1) | 1 | 4 | 2 | 3 | >1 |

| Metric | Rank |  |  |  | FF |
| --- | --- | --- | --- | --- | --- |
|  | M1 | M2 | M3 | M4 |  |
| Microbial peptides identified (MSLab2-SE1) | 1 | 3 | 2 | 4 |  |
| Microbial peptides identified (MSLab1-SE2) | 1 | 4 | 2 | 3 | >1 |
| Microbial peptides shared with FF (MSLab1-SE1) | 1 | 3 | 1 | 4 |  |
| Microbial peptides shared with FF - PSM threshold (MSLab1-SE1) | 1 | 3 | 2 | 4 |  |
| Microbial peptides shared with FF (MSLab1-SE2) | 1 | 3 | 1 | 4 |  |
| Microbial peptides shared with FF - PSM threshold (MSLab1-SE2) | 2 | 3 | 1 | 4 |  |
| Alpha diversity (Shannon's index) | 1 | 4 | 2 | 3 | >3 |
| Beta diversity (distance from FF) | 2 | 1 | 3 | 4 |  |
| Correlation with FF | 3 | 1 | 2 | 4 |  |
| Reproducibility (correlation between replicates) | 1 | 2 | 3 | 4 |  |
| Relative abundance of hydrophobic peptides (distance from FF) | 2 | 3 | 4 | 1 |  |
| Taxonomic annotation yield | 3 | 1 | 4 | 2 | >1 |
| Taxonomic richness | 2 | 3 | 1 | 4 | >3 |
| Taxa shared with FF | 2 | 3 | 1 | 4 |  |
| Taxa shared with FF - PSM threshold | 1 | 3 | 2 | 4 |  |
| Bacillota/Bacteroidota ratio (distance from FF) | 1 | 2 | 3 | 4 |  |
| Functional annotation yield | 3 | 4 | 2 | 1 | >4 |
| Functional richness | 1 | 4 | 2 | 3 | >3 |
| Functions shared with FF | 1 | 3 | 1 | 4 |  |
| Functions shared with FF - PSM threshold | 1 | 3 | 1 | 3 |  |
| Carbon metabolism pathway coverage | 2 | 4 | 1 | 3 | >3 |

\*FF ranking with respect to the four stabilization methods. ">1" and ">3" mean that the FF value is higher than the best-ranked and the third-ranked stabilization methods, respectively.

In general, FF still exhibited the best performance in terms of identification yield and taxonomic annotation yield. The poorer identification rates of stabilized samples may be due to the presence in the preservation media of molecules able to interfere with peptide quantitation prior to LC-MS/MS analysis, with precursor ion selection during MS analysis, or with peptide/protein identification by induction of unexpected chemical modifications. Furthermore, the freeze–thaw cycle applied prior to sample stabilization, as necessary for shipping aliquots between laboratories, constitutes an artificial step that would not typically occur in real-world collection settings. To control for this variable, we designed our study to apply the same number of freeze– thaw cycles (two) to both FF and stabilized samples. Nevertheless, performing a freeze–thaw cycle before stabilization may have further contributed to uneven bacterial cell lysis and/or introduced bias in peptide and protein identification due to protease activation (from host or microbial sources) or irreversible protein aggregation and precipitation upon thawing. Overall, M1 ranked in first place for 19 out of the 28 metrics measured in the study, followed by 8 for M3, 3 for M2, and 2 for M4. Specific strengths shown by M1 were the highest number of distinct peptides identified, the highest alpha diversity, the highest level of reproducibility between replicates, the highest functional richness, and the Bacillota/Bacteroidota ratio closest to the FF control. The main advantages found for M2 were instead the high taxonomic annotation yield, the shortest distance from the FF control in terms of beta diversity, and the highest correlation level of the peptide abundance profiles with the FF control. M3 outperformed the other methods in terms of taxonomic richness and coverage of the carbon metabolism pathway. Finally, M4 stood out for the relative abundance of hydrophobic peptides closest to the FF control and for the functional annotation yield.

Based on the degree of overlap between identifications and on the PCA results, the metaproteome profiles obtained with M1 and M3 (i.e. the methods with the highest identification yields) were globally quite comparable. However, both stabilization methods led to the identification of a considerable fraction of peptides and related functions that were not identified in any of the FF replicates. If an increase in the relative abundance of certain taxa (e.g. Bacillota for M3) can be explained by a lower or higher resistance to the lysis induced by the distinctive components of each method, it is more complex to understand the reason for the significantly higher relative abundance of many specific proteins (including ribosomal proteins, chaperonins, flagellins, tRNA synthetases, and enzymes involved in butyrate biosynthesis) in the samples stabilized with M1 and M3. We could mention two possible hypotheses (not necessarily alternative to each other) that could explain this phenomenon: an increased efficiency of cell lysis related to the long exposure to the detergents contained in the two media, and a rapid activation of some basic biological processes (including replication, transcription, and translation) in the microbial cells as a response to the stabilization environment, before cell lysis occurs. According to the first hypothesis, M1 and M3 would be credited with improving the coverage of the metaproteomic profile by allowing the identification of functions which were actually expressed at the time of sample collection, but whose signal was overwhelmed by that of other more abundant peptides in the FF control. According to the second hypothesis, these stabilization treatments would introduce substantial biases that would lead to significant but artifactual changes in the functional profile of the fecal microbiome compared to the time of collection. However, strong chaotropes such as LDS and guanidine thiocyanate are expected to denature proteins quickly, possibly before microbial cells can orchestrate an appropriate molecular response. Additionally, to the best of our knowledge, no study has investigated the lysis efficiency or protein solubilization properties of the main components of the tested solubilization methods (e.g., guanidine thiocyanate and LDS) toward different bacterial types and their external structures, or even explored the biophysical aspects of the interaction between these molecules and the cell membranes or walls of various bacterial types. In all cases, although metaproteomic analysis should ideally reflect the exact conditions of the sample at the time of collection, any intervention for its preservation or transport (i.e., freezing or other stabilization methods) can potentially introduce artifacts. Nevertheless, it is essential that such artifacts remain as reproducible as possible. Therefore, future studies are needed to address these specific issues and to further validate the reliability, robustness, and unbiasedness of M1 and M3 as stabilization methods for fecal metaproteomics.

Several considerations can be made regarding the advantages and disadvantages of each stabilization method with respect to their potential large-scale application in fecal metaproteomics-enabled population studies. M1 is based on the FDA-cleared, commercially available eNAT^®^ system, which combines a flocked swab for sample collection with a guanidine thiocyanate-based medium for sample preservation. Considering its ease of use and its validation for nucleic acid-based fecal microbiome analysis, this device can be recommended for large studies integrating metaproteomics with metagenomic and/or metatranscriptomic approaches. In our multicenter study, the eNAT^®^-based protocol was also the most robust, showing by far the highest reproducibility between sLab replicates. This is consistent with the fact that this method is the one with the lowest number of steps and the shortest duration. However, in “real world” settings, the use of fecal swabs deserves careful instruction of untrained individuals engaged in self-sampling, as a greater or lesser amount of specimen collected than required may compromise the efficacy of the guanidine thiocyanate-based medium. M2, based on acetone precipitation of proteins, appears to be potentially less manageable when sampling occurs in a hospital or home setting, as it requires more skill and time than the other four methods. M3 is based on the Invitrogen™ LDS sample buffer. This procedure requires sufficient skill to adequately mix the buffer and the specimen, as well as a heater to reach high temperatures for microbial inactivation. It can be adopted when sampling occurs in specific settings, close to a minimally equipped laboratory or a clinical ward. In addition, it has the potential to be developed into a kit for home use. Finally, M4 is built around an RNAlater-like medium. As mentioned above, it was the only one of the five methods analyzed here that had previously been used in metaproteomic studies. When tested with human feces, RNAlater showed fewer protein identifications compared to the FF control, along with a higher relative abundance of Bacteroidetes and other Gram-negative bacteria [25]. However, in their study, Hickl et al. processed the FF samples (but not the RNAlater-treated samples) by cryomilling, which could increase the lysis of Gram-positive bacteria and, in turn, lower the relative abundance of proteins encoded by Gram-negative bacteria, including Bacteroidetes. In our study, RNAlater-treated samples (M4) showed a strikingly different Bacillota to Bacteroidota ratio compared to FF, which appeared to be largely driven by a reduced abundance of *Bacteroides*, typically one of the main contributors to the Bacteroidota phylum in human feces. A possible explanation for this reduction is that *Bacteroides* spp. are distinguished within the Bacteroidales order by their comparatively thick and highly regulated polysaccharide capsules [27,28]. The addition of RNAlater, with its high-salt environment (notably its ∼70% saturated ammonium sulfate content), may promote precipitation or structural alteration of these extracellular polysaccharides, thereby affecting downstream protein solubility and detection. In another study, the RNAlater medium was successfully tested on mouse feces with minimal differences in terms of identification yield compared to the FF control [9]. These two previous studies and the present one cannot be easily compared as the reagents and protocols were slightly different. In addition, mouse and human fecal samples display strikingly different textures, which may affect the quality of protein extraction with this medium. Finally, M5 (OMNImet^®^•GUT device) was excluded from the main analyses because the number of peptides identified was extremely low and incomparable with the other methods. The reason for this poor performance might be related to the high concentration of ethanol (>60%) in the stabilization medium, which was not completely removed before the methanol/chloroform precipitation step. However, the OMNImet^®^•GUT device is of great interest because of its reported ability to preserve fecal (microbial and human) metabolites. Therefore, its potential use in metaproteomics deserves further evaluation through the development of specific experimental procedures aimed at obtaining high-quality protein extracts. In general terms, further optimization of sample preparation protocols tailored to each specific stabilization method, which is beyond the scope of this particular study, is expected to improve the identification yield and the depth of the metaproteome profiles obtained from stabilized fecal samples.

## Conclusions

This collaborative multicenter study evaluated five methods for preserving fecal microbiome samples for metaproteomic analysis. While all stabilization methods exhibited their own strengths and weaknesses, M1 (i.e. the e-NAT^®^ collection and preservation system) appears to hold the greatest potential for advancing metaproteomic research and expanding the knowledge of gut microbiome functions, primarily because of its remarkable reproducibility. Easy-to-use methods are known to reduce operator bias in any given setting, consistent with the need for stabilization procedures to steadily control variability. In this regard, methods such as M3 and M4 have potential to be implemented for broader applications, including the development of user-friendly home kits. Future efforts will allow further optimization of all the methods and improve our knowledge of their interaction with microbial cells, including the mechanisms behind potential biases in the metaproteome profile. Current advancements in fecal sample stabilization are expected to stimulate standardized, large-scale population studies involving metaproteomic analysis and thus significantly accelerate progress in gut microbiome research. Together with the improvements of bioinformatic workflows currently under study within the Metaproteomic Initiative framework (https://metaproteomics.org/campi/campi3), these important efforts will be key to further strengthen the robustness of research methods in metaproteomics.

## Methods

### Sample collection and aliquoting

A fecal sample (approximately 180 g) was provided by a healthy 54-year-old female subject. The donor signed a formal written consent form, providing permission for the analysis of her fecal samples and the publication of the related case study (study protocol “PATH_TCA”, approved by the Ethical Committee of AOU Cagliari, Italy). The sample was collected in a sterile bag, cooled with an ice pack and delivered in 30 minutes by a healthcare professional to the aLab (Uzzau Lab, Department of Biomedical Sciences, University of Sassari, Italy). The sample was immediately manually homogenized inside the bag for 15 min (kept on ice). The bag was opened lengthwise under a biological hood and placed on a bed of ice for aliquoting. Within two hours, the fecal sample was divided into aliquots of approximately 3 g each, which were placed in previously autoclaved and weighed 5 mL tubes, kept on ice until aliquoting was complete. One aliquot was placed in a standard stool container and immediately subjected to standard bacterial culture tests that confirmed the absence of enteric pathogens, while the other aliquots were immediately frozen at −80°C.

Within one month, sixteen frozen stool aliquots were shipped on dry ice to the eight sLabs involved in the study (two aliquots per sLab). An additional five aliquots were thawed at 20°C until melting, subjected to sample stabilization according to the five methods described below, and then subjected to bacterial culture tests to assess bacterial viability. No viable bacteria were detected in any of the stabilized samples. A final frozen stool aliquot was shipped on dry ice directly from the aLab to the pLab for use as a flash-frozen (FF) control.

### Sample stabilization

Sample stabilization was performed in parallel in eight different sLabs: Armengaud Lab, Li2D, CEA, Bagnols-sur-Cèze, France; Bastida Lab, Department of Soil and Water Conservation and Organic Waste Management, CEBAS-CSIC, Murcia, Spain; Benndorf Lab, Bioprocess Engineering, Otto von Guericke University, Magdeburg, Germany; Hernandez-Raquet Lab, Toulouse Biotechnology Institute – INRAE, University of Toulouse, Toulouse, France; Jehmlich Lab, Department of Molecular Toxicology, Helmholtz-Centre for Environmental Research-UFZ, Leipzig, Germany; Seifert Lab, Institute of Animal Science, University of Hohenheim, Stuttgart, Germany; VIB Proteomics Core, VIB – UGent Center for Medical Biotechnology, Ghent, Belgium; Wilmes Lab, Luxembourg Centre for Systems Biomedicine, University of Luxembourg, Esch-sur-Alzette, Luxembourg.

Upon delivery, each sLab checked that the stool aliquots were still frozen and immediately stored them at −80°C. On the day scheduled for sample stabilization in all sLabs, the stool aliquots were thawed in a water bath at 20°C until melting, placed in an ice bucket under a sterile hood, and subjected to sample stabilization using the five selected methods (M1-M5) according to the protocols described below.

Method 1 (M1) was based on the e-NAT^®^ collection and preservation system (6U073S01; Copan, Brescia, Italy). Approximately 200 mg of stool sample was collected using the swab provided with the device. The swab was then placed in the provided tube containing the stabilizing medium. According to the manufacturer, the e-NAT stabilizing medium contains guanidine thiocyanate, Tris-EDTA, HEPES, and an unspecified detergent. The tube was vortexed for 1 min to thoroughly dissolve the sample.

Method 2 (M2) was acetone based. Approximately 200 mg of stool sample was transferred to a safe-lock tube. Then 2 ml of pre-chilled acetone was added to the tube. The tube was vortexed for 2 min to thoroughly dissolve the sample and incubated for 1 hour at –20°C. After that, the sample was centrifuged at 10,000×*g* for 10 min at 4°C and allowed to dry overnight at room temperature until complete acetone evaporation.

Method 3 (M3) was based on the LDS sample buffer (Invitrogen™, Thermo Fisher Scientific, Waltham, USA). Approximately 200 mg of stool sample was transferred to a safe-lock tube. Then 200 μL LDS 4X sample buffer and 400 μL MilliQ water were added sequentially to the tube. The tube was vortexed for at least 3 min to homogenize the sample and then heated for 5 min at 99°C to inactivate the sample.

Method 4 (M4) was based on an RNAlater-like preservation buffer prepared as follows: 40 ml of 0.5 M EDTA, 25 ml of 1 M trisodium citrate dihydrate and 700 g of ammonium sulfate were added sequentially to 935 ml of DEPC water at 50°C. After cooling, the pH was adjusted to 5.2 with sulfuric acid. Approximately 200 mg of stool sample was transferred to a safe-lock tube. Then, 200 μL RNAlater-like buffer was added to the tube. The tube was vortexed for at least 3 min to thoroughly dissolve the sample.

Method 5 (M5) was based on the OMNImet^®^•GUT device (DNA Genotek, Stittsville, Canada). Approximately 500 mg of stool sample was collected using the spatula provided with the device and transferred into the tube top. After the cap was screwed back on the tube top, the sealed tube was shaken vigorously for at least 30 sec to dissolve the sample.

All stabilized samples were stored at room temperature for 14 days and then shipped to the pLab (Armengaud Lab, Li2D, CEA, Bagnols-sur-Cèze, France). Upon arrival at the pLab, stabilized samples were immediately stored at –20°C until protein extraction.

The FF control was prepared at the pLab as follows: after thawing in a water bath at 20°C, approximately 200 mg of fecal material was resuspended in 400 μL of LDS 1X sample buffer (Invitrogen™, Thermo Fisher Scientific) and immediately stored at –20°C until protein extraction.

### Protein extraction and digestion

Protein extraction and digestion from all stabilized and FF fecal samples was performed at the pLab. Samples were transferred to 2-ml screw cap microtubes (Sarstedt, Marnay, France) containing 200 mg of beads, as previously described [12]. Samples were sonicated for 5 min in an ultrasonic water bath (VWR ultrasonic cleaner). Cell disruption was performed on a Precellys Evolution instrument (Bertin Technologies, Aix-en-Provence, France) operated at 10,000 rpm for 10 30-s cycles, with 30 s rest between cycles. Following lysis, samples were centrifuged at 16,000×*g* for 3 min.

For samples stabilized using Methods 1, 4 and 5, proteins in the supernatant were precipitated with the sequential addition (3:4:1 ratio) of ice-cold water (600 µL), methanol (400 μL), and chloroform (100 μL), and vortexed after the addition of all solvents. Samples were centrifuged at 16,000×*g* for 2 min at room temperature. The methanol/water layer was removed from samples (top layer) without disrupting the pellet.

Samples were washed with the addition of 800 μL of ice-cold methanol before being vigorously vortexed and centrifuged at 16,000×*g* for 2 min. The resulting pellet was resuspended in LDS 1X sample buffer (Invitrogen™, Thermo Fisher Scientific). All the samples were incubated at 99°C for 5 min before an aliquot of 25 μL was loaded onto a NuPAGE 4%–12% Bis-Tris gel. Proteins were subjected to short (5-min) SDS-PAGE migration, followed by staining with Coomassie SimplyBlue SafeStain (Thermo Fisher Scientific) for 5 min. In-gel trypsin proteolysis was then performed using Trypsin Gold (Promega) using 0.011% ProteaseMAX surfactant (Promega, Madison, WI, USA), as described by Hartmann et al. [29]. The resulting peptide pools were dried in MS vials using a SpeedVac and then transferred at ambient temperature to the two MS labs (listed below) for metaproteome measurements.

### LC-MS/MS analysis

Peptide mixtures were analyzed in duplicate at the MSLab1 (ProGénoMix platform, CEA, France) and at the MSLab2 (ORNL, USA).

At MSLab1, peptide mixtures were reconstituted with 45 μL of 0.1% trifluoroacetic acid and quantified using the Pierce™ Quantitative Peptide Assays & Standards (Thermo Fisher Scientific). After randomization, a 200-ng aliquot of each peptide mixture was injected and analyzed on a Q Exactive HF tandem mass spectrometer (Thermo Fisher Scientific) coupled to an Ultimate 3000 Nano LC System (Thermo Fisher Scientific). Peptides were injected onto a reverse-phase on a Acclaim PepMap100 C18 pre-column (5 μm, 100 Å, 300 μm id × 5 mm at 45°C) and a Acclaim PepMap 100 Neo C18 column (2 μm, 100 Å, 75 μm id × 500 mm), desalted, and then resolved at a flow rate of 0.2 μL/min with a 120 min gradient of acetonitrile (4-20% for 100 min, then 20-32% for 20 min) in the presence of 0.1% formic acid. The tandem mass spectrometer operated in data-dependent mode, using a Top20 strategy, selecting parent ions with 2+ or 3+ charge states and a dynamic exclusion window of 10 sec. Full-scan mass spectra were acquired from 350 to 1,500 m/z at a resolution of 60,000 with internal calibration activated on the m/z 445.12002 signal. MS/MS scans were acquired at a resolution of 15,000 with an intensity threshold of 1.7 × 10^4^ and an isolation window of 0.7 m/z.

At MSLab2, peptide mixtures were reconstituted with 50 μL of 0.1% trifluoroacetic acid and quantified using a Nanodrop Scopes A205. After randomization, a 2 ug aliquot of each peptide sample in solution was analyzed by automated 1D LC-MS/MS analysis using a Vanquish ultra-HPLC (UHPLC) system plumbed directly in-line with a QExactive-Plus mass spectrometer (Thermo Fisher Scientific). An in-house-pulled 100 µm inner diameter nanospray emitter was packed to 15 cm with 1.7 µm Kinetex C18 reverse-phase resin (Phenomenex). Peptides were loaded and separated by uHPLC (at ambient temperature) under the following conditions: sample direct load injection followed by 100% solvent A2 (98% water, 2% acetonitrile, 0.1% formic acid) from 0 to 30 min to load, a linear gradient from 0 to 30% solvent B (70% acetonitrile, 30% water, 0.1% formic acid) from 30 to 220 min for separation, and 100% solvent A from 220 to 275 min for column re-equilibration. Eluting peptides were directed into the nano-electrospray source (operated at a flow rate of 0.2 μL/min) and analyzed with the following MS settings: data-dependent acquisition, top-10 method; mass range 300–1500 m/z; MS and MS/MS resolution 70 and 15 K, respectively; MS/MS loop count 20; isolation window 1.8 m/z; charge state exclusion of unassigned, +1, +6–8 charges.

### Generation of a matched metagenomic database

DNA extraction, preparation, and sequencing were performed at the aLab. DNA was extracted from two fecal sample aliquots with the QIAamp Fast DNA Stool Mini Kit (Qiagen Hilden, Germany) to perform a shotgun sequencing of the whole fecal metagenome. DNA extracts were quantified using a Qubit™ 4 Fluorometer with the dsDNA High Sensitivity assay kit (Thermo Fisher Scientific, Waltham, MA, United States) and then diluted to 4 ng/μl. The DNA was subjected to tagmentation and ligation of MiSeq adaptors according to the instructions of the Nextera XT kit (Illumina, San Diego, CA, United States). Libraries (average size of 464 bps) were validated using the Agilent 4150 TapeStation system with the D1000 ScreenTape assay kit (Agilent Technologies, Santa Clara, CA, United States), quantified with the Qubit dsDNA High Sensitivity assay kit, and finally diluted to equimolar concentrations following Illumina guidelines. Libraries were sequenced using a MiSeq sequencer (Illumina). The MiSeq Reagent Kit v3 from Illumina was used (following the manufacturer’s specifications) to generate paired-end reads of 301 bases in length in each direction. A total of 16,292,837 and 3,310,285 read pairs were obtained from the two fecal sample aliquots sequenced.

Raw reads were processed using a suite of tools available in the Galaxy platform (https://proteomics.usegalaxy.eu). First, reads were trimmed and filtered using Trimmomatic (Galaxy version 0.38.1) [30], applying the ILLUMINACLIP, TRAILING, MINLEN and LEADING steps, all with default parameters (except for MINLEN, which was set to 36). Human sequences were aligned and removed using Bowtie 2 (Galaxy version 2.5.0) [31], with the *Homo sapiens* database “hg38 full” as reference genome and default settings. Reads were then assembled into contigs using MEGAHIT (Galaxy version 1.2.9) [32] with default parameters. Open reading frames were found and translated into amino acid sequences with FragGeneScan (Galaxy version 1.30.0) [33], using “illumina_5” as a model. After removing redundant amino acid sequences using CD-HIT (Galaxy Version 4.8.1) [34] with a sequence identity threshold of 1, the remaining sequences were used as an additional database for peptide identification (see below).

### Bioinformatic analyses

Protein identification was performed using two different bioinformatic pipelines. In both cases, searches were performed against a combination of three sequence databases: the IGC collection of human gut metagenomes (https://ftp.cngb.org/pub/SciRAID/Microbiome/humanGut_9.9M/GeneCatalog/IGC.pep.gz) [35], a matched metagenome obtained from the same fecal sample analyzed by metaproteomics (described in the “Generation of a matched metagenomic database” section), and the *Homo sapiens* reference proteome retrieved from UniProtKB/Swiss-Prot (release 2021_04).

With respect to the first pipeline, the MS raw files were analyzed using the Proteome Discoverer™ software (v.2.5; Thermo Fisher Scientific), with Sequest-HT as the search engine and Percolator for peptide validation, setting the FDR threshold to 1%. The search parameters were as follows: precursor mass range, 350-5000 Da; minimum peak count, 6; S/N threshold, 2; enzyme, trypsin (full); maximum missed cleavage sites, 2; peptide length range, 5-50 amino acids; precursor mass tolerance, 5 ppm; fragment mass tolerance, 0.02 Da; static modification, cysteine carbamidomethylation; dynamic modification, methionine oxidation. Offline mass recalibration and MS1-based label-free quantification (LFQ) were carried out using the Spectrum Files RC and the Minora Feature Detector nodes, respectively [36]. The optimal settings for retention time and mass tolerance windows were calculated by the Minora algorithm based on mass accuracy and retention time variance distribution. A consensus feature list was defined based on the outputs of the Feature Mapper and Precursor Ions Quantifier nodes. MS1 signals of all peptides exhibiting significant matches with at least one MS2 spectrum from at least one sample were mapped across runs and quantified by calculating the integrated area of the chromatographic peak. LFQ data were normalized by selecting the “Total Peptide Amount” normalization mode. Human proteins were grouped based on the maximum parsimony principle and only master proteins (i.e. the most representative member of each protein group) were considered for the analysis. With respect to the second pipeline, MS raw files were converted to mgf files with MS Convert (version 3.0, 32-bit, zlib compression, peak count threshold of 200 most intense peaks). Then, mgf files were submitted for database searches to Mascot Server (version 2.8) using the following search parameters: precursor mass tolerance, 10 ppm; fragment mass tolerance, 20 ppm; #C13, 1; maximum missed cleavages, 2; fixed modification, cysteine carbamidomethylation; variable modification, methionine oxidation; precursor charges, 2+, 3+ and 4+. Searches against decoy databases were also performed. Mascot dat files were used to create a peptide matrix including all PSMs but based on only those proteins that passed the FDR threshold (1%).

Peptide taxonomic annotation was carried out using Unipept Desktop (v.2.0.0) [37], selecting the three available options (“equate I and L”, “filter duplicate peptides” and “advanced missed cleavage handling”). Protein sequences were subjected to functional annotation using eggNOG-mapper (v.2.1.8, available in the Galaxy platform at https://proteomics.usegalaxy.eu) [38] keeping default parameters, and then choosing COG [39] and KEGG orthology (KO) information as main functional classifications [40]. Meta4P (v.1.5.5) was employed to parse identification, quantification and annotation data, as well as to calculate the related metrics [41]. Peptide sequences matching the first and/or the second of the sequence databases listed above (IGC and matched human fecal metagenomes, respectively), but not the *H. sapiens* database, were considered microbial and their LFQ values were renormalized (i.e., each LFQ value was divided by the sum of all microbial LFQ values measured in that sample and multiplied by 10^10^) to approximate their relative abundance within the microbiota. The abundance of a taxon or a function was estimated by summing the LFQ values associated with all peptides having that feature among their annotations. The grand average of hydropathy (GRAVY) score was calculated for each peptide sequence identified in the study using the online GRAVY calculator (https://www.gravy-calculator.de).

### Statistical analysis and graph generation

Correlation analyses were conducted using the R packages *Hmisc* and *DescTools*. Scatter, bar and violin plots were created using GraphPad Prism (v.9.0.0). UpSet plots were generated online using SRplot (www.bioinformatics.com.cn/srplot) [42]. Principal component analysis (PCA) was carried out using the ClustVis web application (https://biit.cs.ut.ee/clustvis) [43], using the following pre-processing options: ln(x+1) transformation, row centering, unit variance scaling, SVD with imputation as PCA method. Heatmaps were generated using Morpheus (https://software.broadinstitute.org/morpheus).

## Additional files

**Additional file 1 (pdf): Supplementary Figures**. Figures S1-S13.

**Additional file 2 (xls): Supplementary Data 1. Relative abundances and log ratios of microbial KO functions**.

## Declarations

### Ethics approval and consent to participate

The donor of the fecal sample analyzed in this study signed a formal written consent form, providing permission for the analysis of her fecal samples and the publication of the related case study (study protocol “PATH_TCA”, approved by the Ethical Committee of AOU Cagliari, Italy).

## Consent for publication

Not applicable.

## Availability of data and materials

Mass spectrometry proteomics data have been deposited to the ProteomeXchange Consortium via the PRIDE [44] partner repository with the dataset identifier PXD060335.

## Authors’ contributions

A.T. contributed as the main author to metagenome sequence processing to generate the custom metagenomic database, to mass spectrometry data processing, annotation, analysis, and visualization, and to manuscript writing and editing. K.S. contributed to mass spectrometry data processing and analysis. S.U. conceived and supervised the multi-laboratory comparison launched in the context of the Metaproteomic Initiative, supervised distribution of fecal samples and reagents, and nucleotide sequencing, conceived and supervised manuscript writing and editing. R.L.H. and J.A. jointly supervised the mass spectrometry analyses, contributed to the data selection, concept, and structure of the manuscript. S.U. and R.H. jointly supervised data analysis and visualization. N.J. and D.B. contributed to the data selection, concept, and structure of the manuscript. M.A., L.D.D., and M.A.D. did DNA extraction, preparation, and sequencing for the custom metagenomic database. L.G., S-B.H., G.M., J.S.S., M.W., F.B., S.D., G.H-R., and B.J.K. did laboratory work for one of the sLabs, which included sample stabilization and stabilized sample distribution. L.G., G.M., and S.L.P. did laboratory work for one of the MS facilities, including protein extraction and proteolytic digestion, and mass spectrometry measurements. J.A., R.L.H., P.W., L.G., R.H., S-B.H., N.J., J.S., D.B., S.D. G.H-R, T.V.D.B., and B.J.K. reviewed and edited the manuscript and participated in discussions on the course of project planning.

## List of abbreviations

aLab: Aliquoting lab
CAMPI: Critical assessment of metaproteome investigation
COG: Cluster of orthologous group
FDR: False discovery rate
FF: Flash-frozen
GRAVY: Grand average of hydropathy
KEGG: Kyoto encyclopedia of genes and genomes
KO: KEGG orthology
LDS: Lithium dodecyl sulfate
LFQ: Label-free quantification
MS: Mass spectrometry
PCA: Principal component analysis
pLab: Preparation lab
SDS: Sodium dodecyl sulfate
SE: Search engine
SIHUMIx: Simplified human gut microbiota model sLab Stabilization lab

## Acknowledgements

This work has benefited from collaborations facilitated by the Metaproteomics Initiative (https://metaproteomics.org/) whose goals are to promote, improve and standardize metaproteomics [2]. NJ thanks Kathleen Eismann for her assistance in the lab.

