## Supplementary Figures for "Critical Assessment of MetaProteome Investigation 2 (CAMPI-2): Multi-laboratory assessment of sample processing methods to stabilize fecal microbiome for functional analysis"

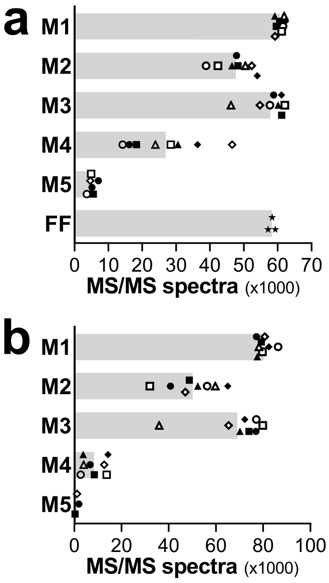


**Figure S1**. Scatter bar plots showing the number of MS/MS spectra acquired at the MSLab1 (**a**) and MSLab2 (**b**) from the same fecal sample treated with the five stabilization methods under comparison (M1-M5) or flash frozen (FF). Each shape marks a different sLab replicate, while the grey bars indicate the mean value for each method.

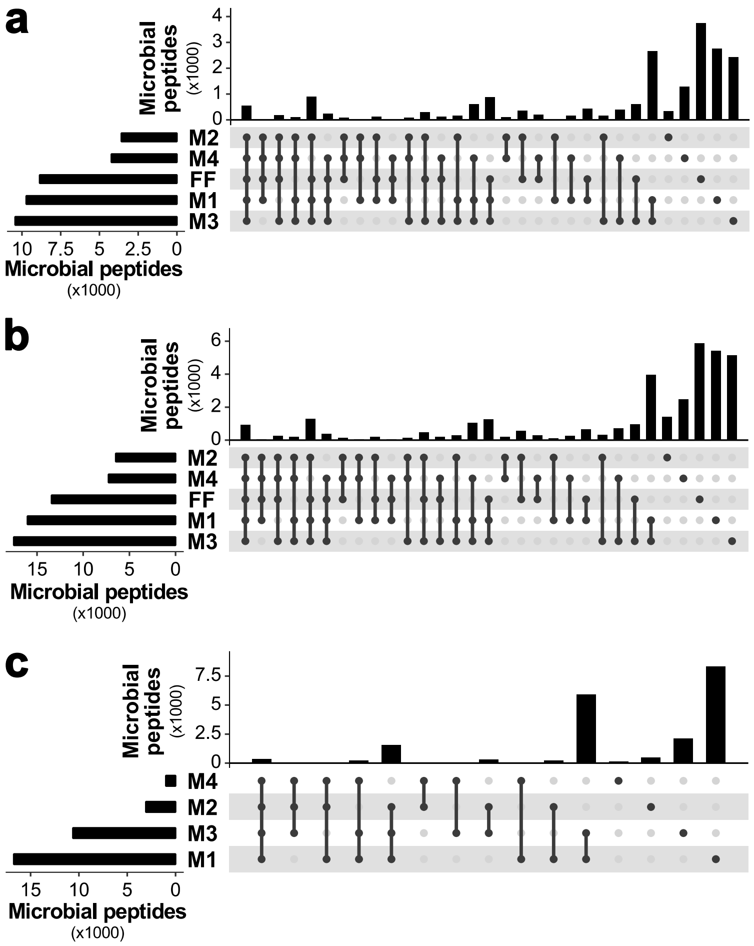


**Figure S2**. UpSet plots showing the microbial peptide identification overlap between the different methods under comparison, according to the MSLab1-SE1 (**a**), MSLab1-SE2 (**b**), and MSLab2-SE1 (**c**) datasets. Peptides identified in the different replicates were merged for each method. Horizontal bars indicate the set size, i.e. the total number of peptides identified for each method (sorted by increasing set size from top to bottom). Vertical bars illustrate the intersection size, i.e. the number of peptide identifications shared between the different methods (sorted by decreasing degree of overlap from left to right), according to the black dot(s) below.

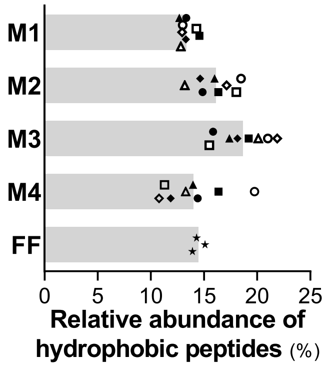


**Figure S3**. Scatter bar plot showing the relative abundance of peptides with a GRAVY score greater than 0.5 (i.e. hydrophobic peptides) in the same fecal sample after processing with four stabilization methods (M1-M4) or flash freezing (FF), according to LFQ data from the MSLab1-SE1 dataset. Each shape marks a different sLab replicate, while the grey bars indicate the mean value for each method.

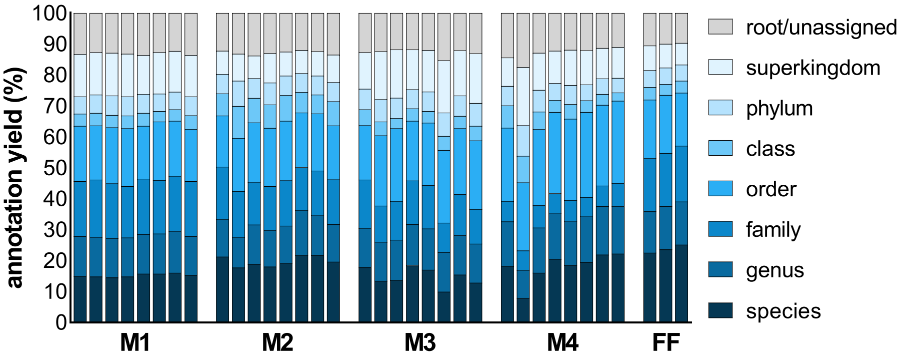


**Figure S4**. Annotation yield at different taxonomic levels obtained with the different sample groups compared in the study, based on the peptide identification results from the MSLab1-SE1 dataset. Each column represents a different sLab (M1-M4) or run (FF) replicate.

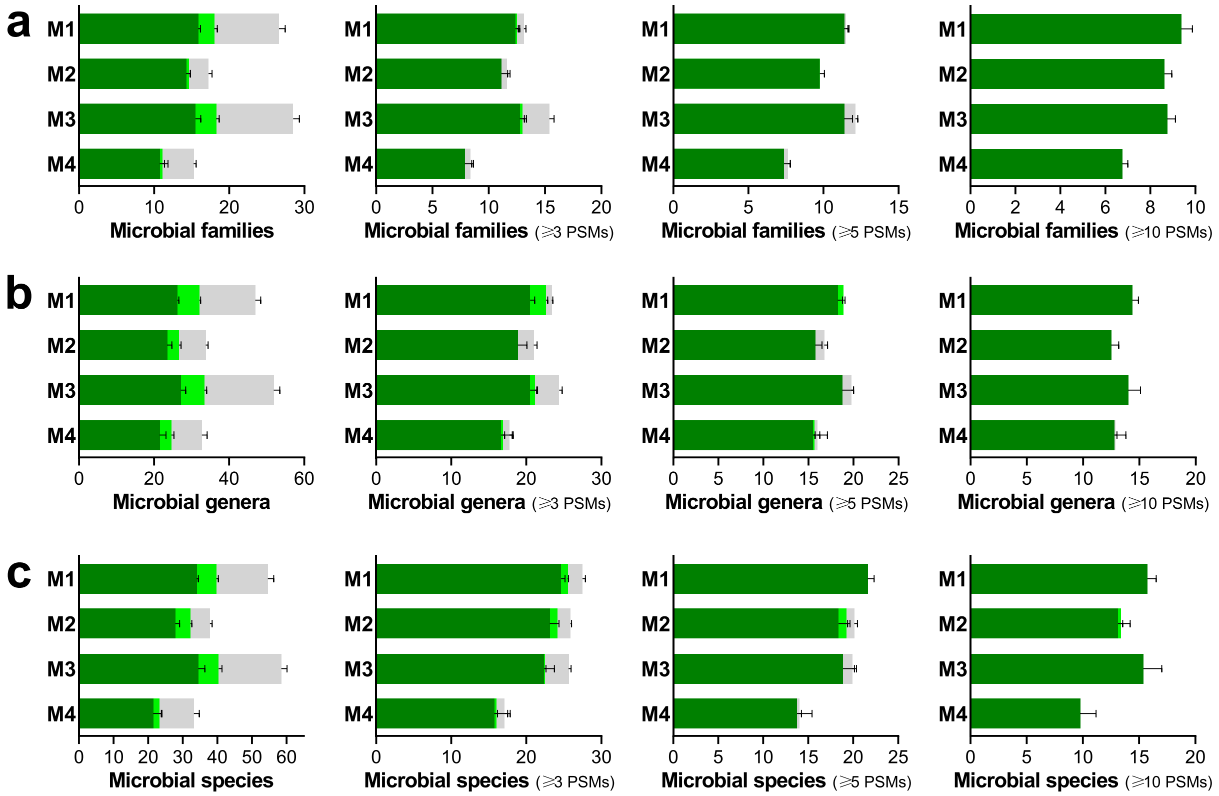


**Figure S5**. Overlap of the identified microbial families (**a**), genera (**b**), and species (**c**) between the four stabilization methods and the FF control. Each bar indicates the mean number of families/genera/species identified between sLab replicates, with error bars indicating the standard error of the mean. Bars are divided into three areas based on the degree of overlap with the FF control: the dark green areas include families/genera/species that were also identified in all three FF replicates, the light green areas include families/genera/species that were also identified in 1 or 2 FF replicates, and the gray areas include families/genera/species that were not identified in any FF replicate. The left plot refers to all the families/genera/species identified, while the others were generated after setting increasing thresholds based on the minimum number of PSMs detected per family/genus/species (namely 3, 5, and 10).

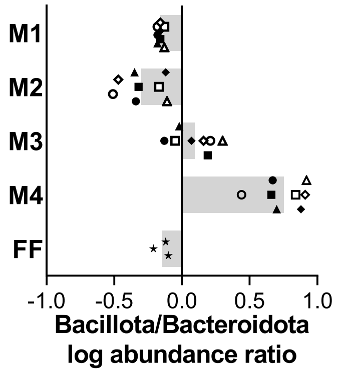


**Figure S6**. Scatter bar plot showing the log ratios between the relative abundances of Bacillota and Bacteroidota, as measured for the four stabilization methods and the FF control in the MSLab1-SE1 dataset. Abundances were calculated by summing LFQ values of all peptides unambiguously assigned to the phyla Bacillota and Bacteroidota, respectively. Each shape marks a different sLab replicate, while the grey bars indicate the mean value for each method.

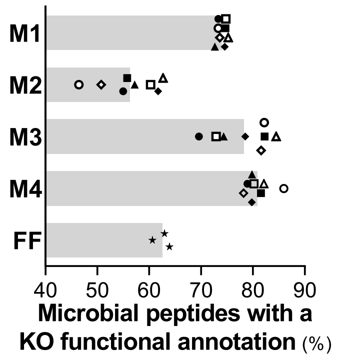


**Figure S7.** Functional annotation yield, measured as the percentage of microbial peptides identified and assigned a KEGG Orthology (KO) annotation, based on the MSLab1-SE1 dataset. Each shape marks a different sLab replicate, while the grey bars indicate the mean value for each method.

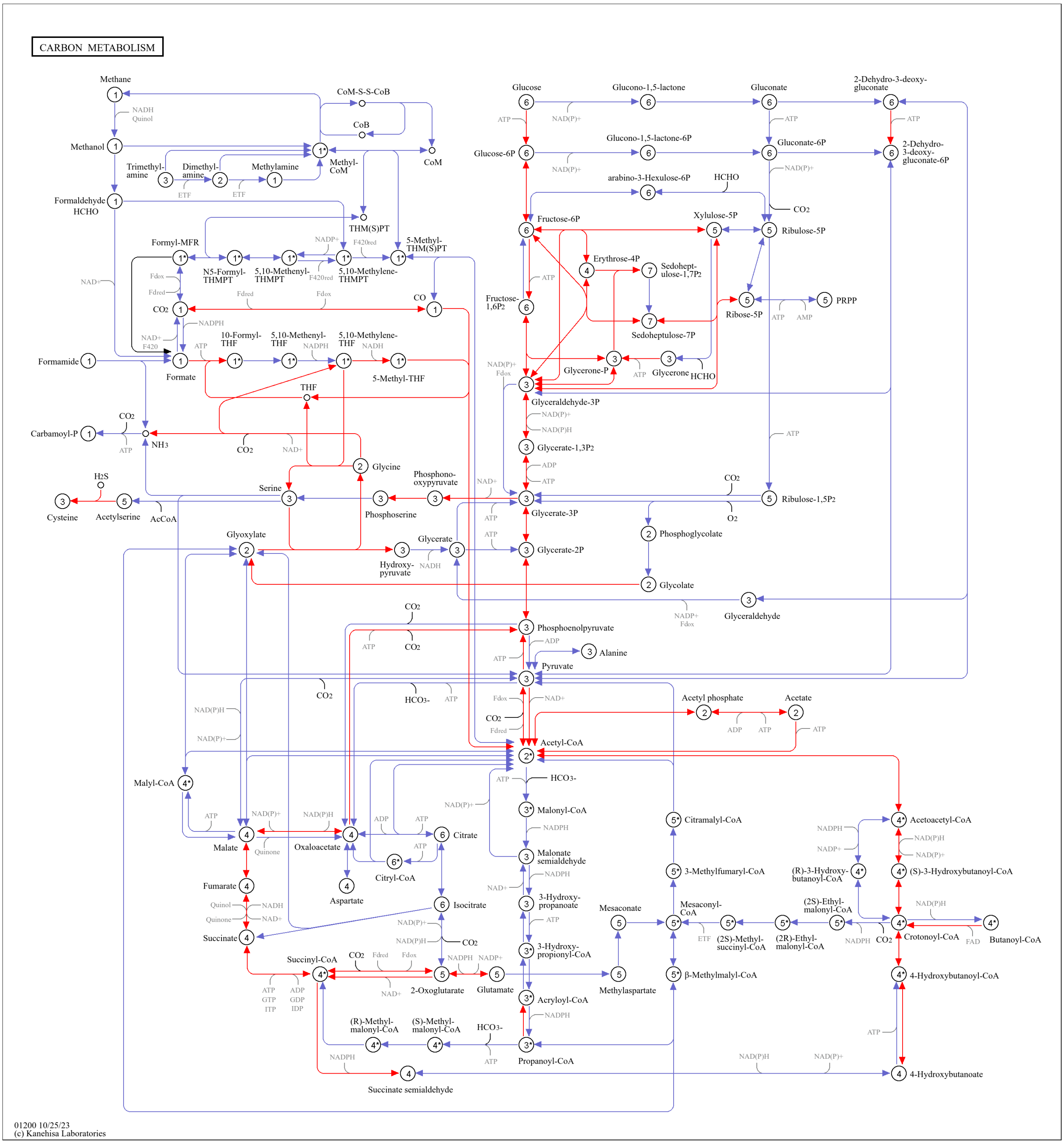


**Figure S8**. KEGG Carbon metabolism pathway map, with microbial KO functions identified in at least two FF replicates (according to the MSLab1-SE1 dataset) colored in red.

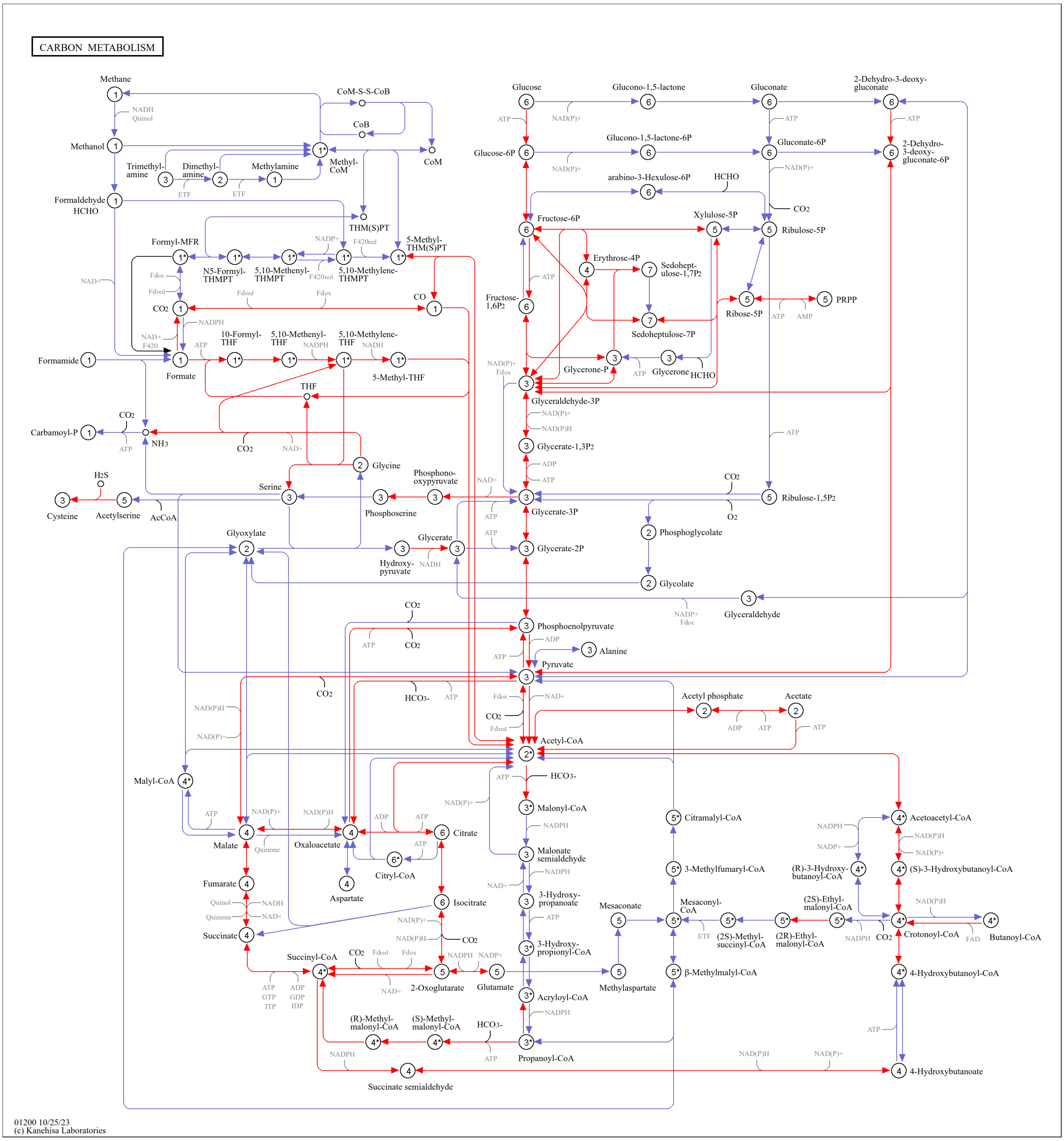


**Figure S9**. KEGG Carbon metabolism pathway map, with microbial KO functions identified in at least two M1 replicates (according to the MSLab1-SE1 dataset) colored in red.

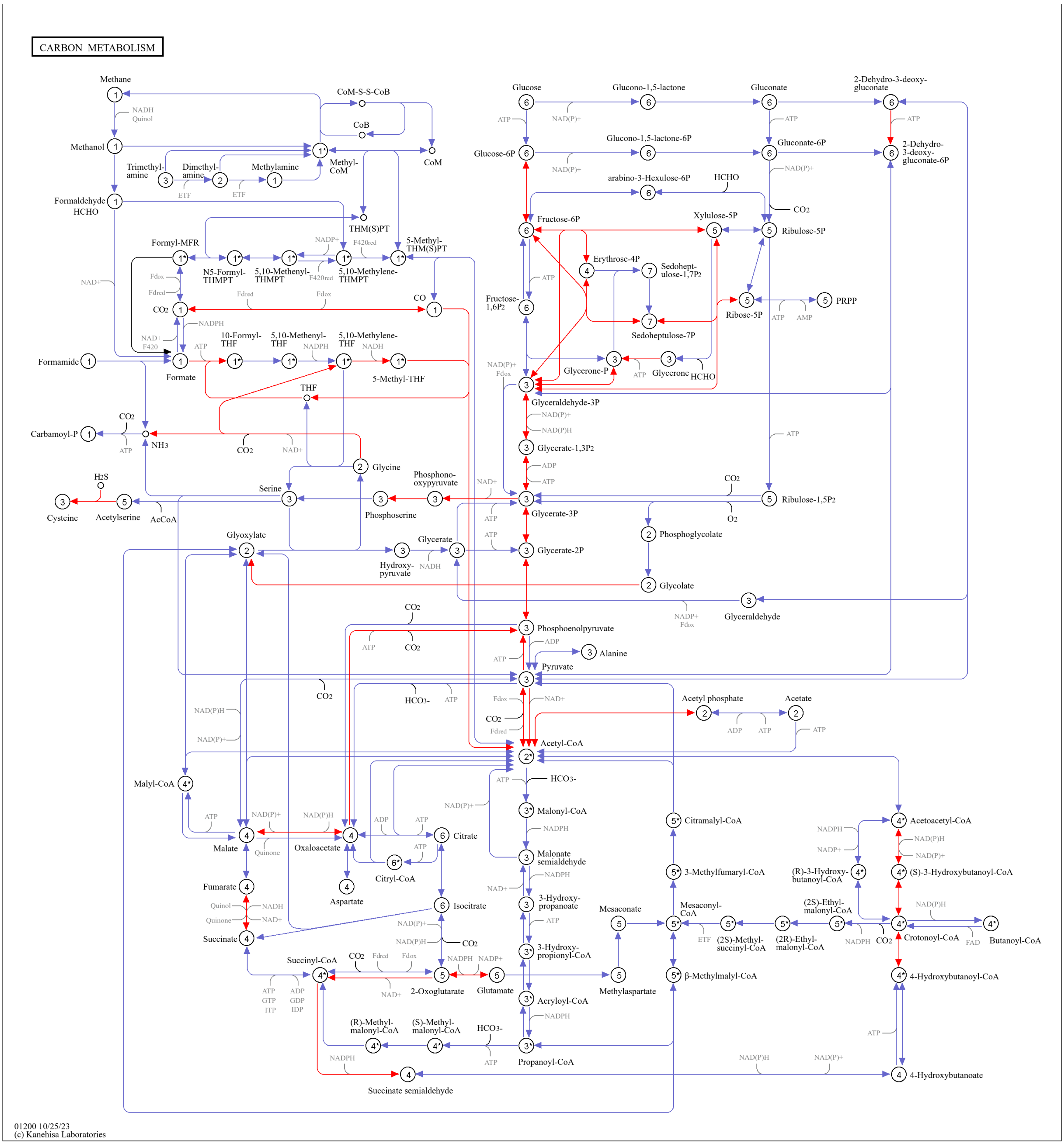


**Figure S10**. KEGG Carbon metabolism pathway map, with microbial KO functions identified in at least two M2 replicates (according to the MSLab1-SE1 dataset) colored in red.

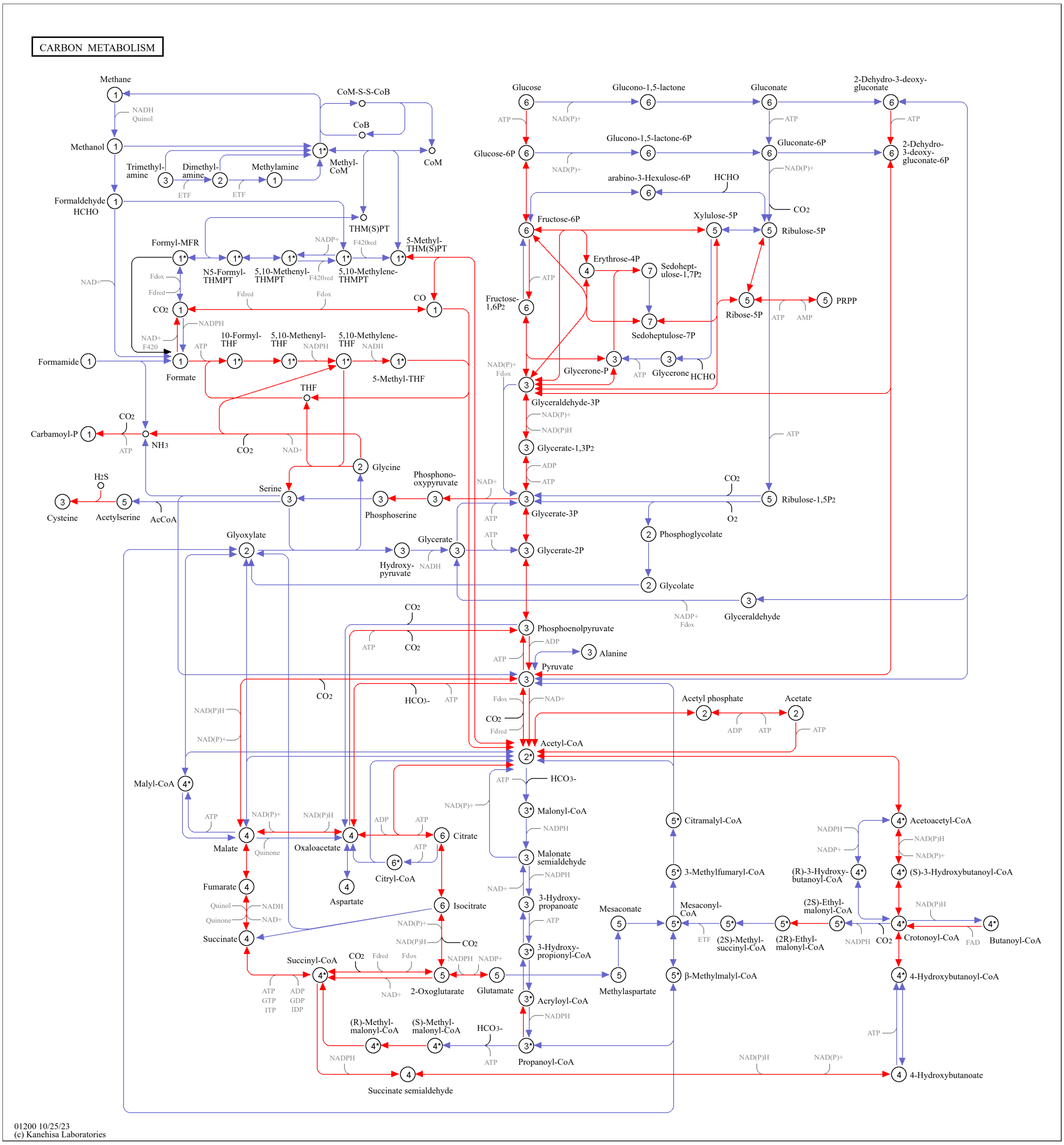


**Figure S11**. KEGG Carbon metabolism pathway map, with microbial KO functions identified in at least two M3 replicates (according to the MSLab1-SE1 dataset) colored in red.

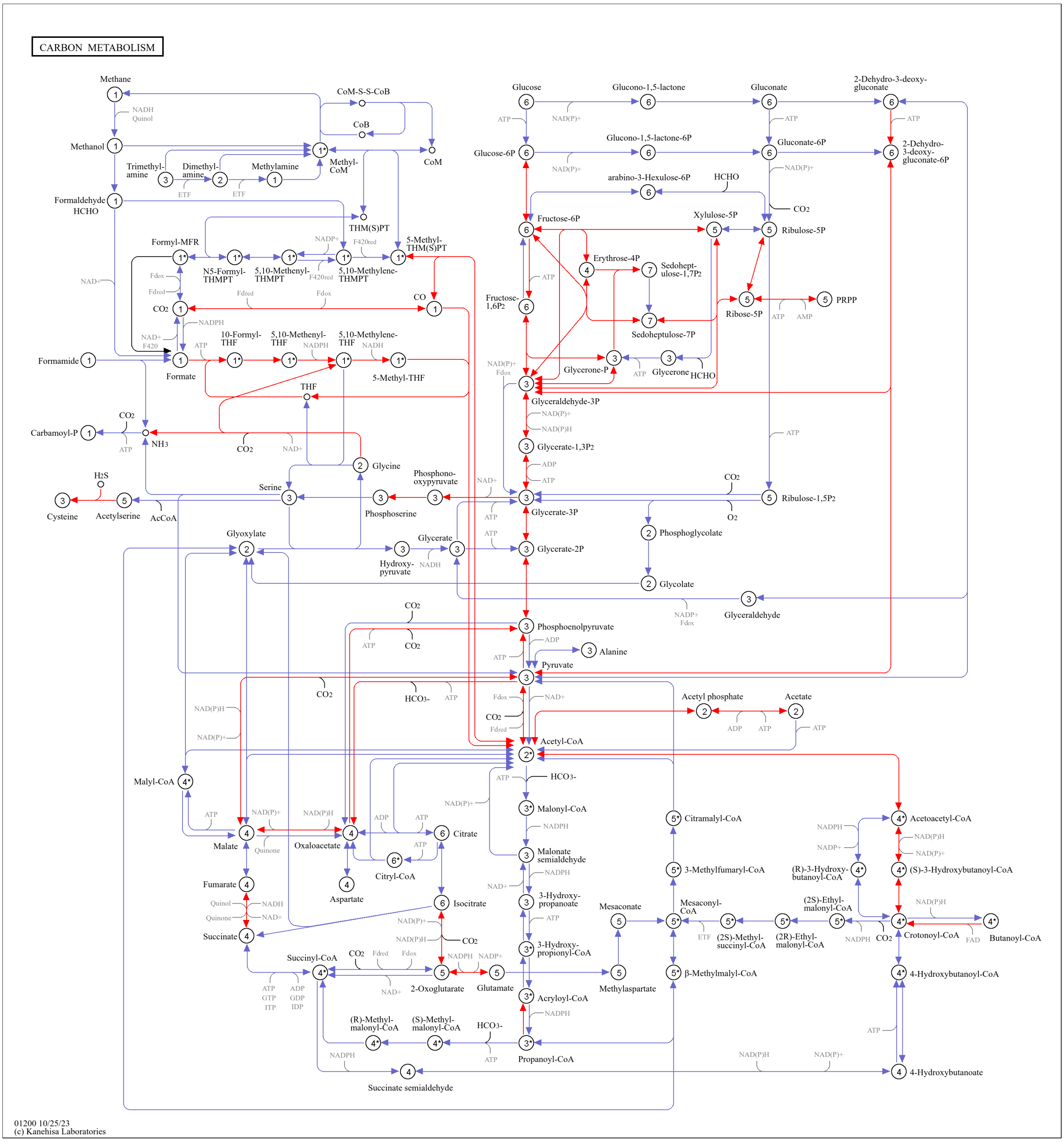


**Figure S12**. KEGG Carbon metabolism pathway map, with microbial KO functions identified in at least two M4 replicates (according to the MSLab1-SE1 dataset) colored in red.

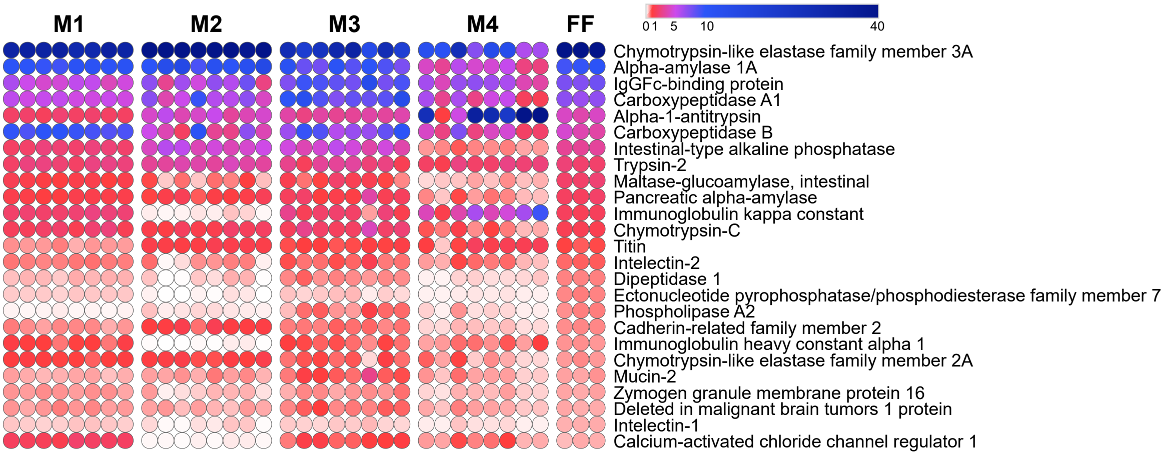


**Figure S13**. Relative abundances of the top 25 human proteins, selected based on their mean relative abundance in the FF group, according to LFQ data from the MSLab1-SE1 dataset. Abundance values are expressed as a color gradient according to the legend in the upper right corner of each heatmap. Each dot represents a different sample replicate.
